# Modest transcriptomic response to polyploidization in allohexaploid wheat synthetics

**DOI:** 10.1101/2022.11.04.515153

**Authors:** Meriem Banouh, David Armisen, Annaig Bouguennec, Cecile Huneau, Mamadou Dia Sow, Caroline Pont, Jerome Salse, Peter Civan

## Abstract

Bread wheat is a recent allohexaploid (genomic constitution AABBDD) that emerged through a hybridization between tetraploid *Triticum turgidum* (AABB) and diploid *Aegilops tauschii* (DD) <10,000 years ago. The hexaploidization can be re-created artificially, producing synthetic wheat that has been used to study immediate genomic responses to polyploidization. Here we produced several synthetic wheats from alternative parental genotypes and reciprocal crosses, and examined transcriptomes from two different tissues and successive generations. We did not detect a massive reprogramming in gene expression, with only ∼1% of expressed genes showing significant differences compared to their lower-ploidy parents. Most of the differential expression is located on the D subgenome, without consistency in the direction of the expression change. Leaves and developing endosperm show distinct patterns of homoeologous expression bias, and almost non-overlapping sets of differentially expressed genes, implying that the polyploidization-triggered reprogramming is not effectuated through permanent (epi)genetic changes. While 0-3 families of transposable elements (TEs) became upregulated in wheat synthetics, we did not detect any significant association between TEs and the expression of nearby genes. We conclude that the modest tissue-specific and partially genotype-specific transcriptomic response to polyploidization is likely caused by rare incompatibilities of parental regulomes, and we discuss the pitfalls of transcriptomic comparisons across ploidy levels that can inflate the de-regulation signal.

## Introduction

Bread wheat (*Triticum aestivum* L.) is one of the world’s crucial staple crops (Shewry and Hey 2015; https://www.fao.org/faostat/en/#data). Considerable efforts are being dedicated to understand its genetic organization, diversity and environmental adaptations, paving the way towards improved performance through genome-informed breeding. Bread wheat is an allohexaploid (2n = 6x = 42 chromosomes; genomic constitution AABBDD) that emerged through a hybridization between allotetraploid wheat *T. turgidum* (AABB) and *Aegilops tauschii* (DD), and subsequent whole-genome duplication (McFadden and Sears 1946). Since the donor of the AABB genomes appears to be a free-threshing, durum-like wheat (Zhou et al. 2020; Zhao et al. 2022) and the major contributor of the D genome is currently distributed along the south-western shores of the Caspian see (Gaurav et al. 2021), wheat allohexaploidization likely occurred after the domestication and spread of free-threshing tetraploids, i.e. <10,000 years ago.

Thanks to the allohexaploid nature of bread wheat, most genes have corresponding copies (homoeologs) originating from the three parental subgenomes. This pool of variably diverged gene variants is expected to provide a heterosis effect that could be behind the crop’s increased yield and adaptive plasticity (Chen 2010; Salman-Minkov et al. 2016). However, transcriptomic studies across several tissues and developmental stages in wheat (Ramirez-Gonzales et al. 2018; Mutti et al. 2018) have revealed that 21.1%-37.4% of triads (homoeologous groups with one copy on each of the three subgenomes) have one or two gene copies silenced. This homoeolog expression bias is in most cases consistent across different tissues (Ramirez-Gonzales et al. 2018), meaning that many genes appear to be permanently silenced in allohexaploid wheat.

The above observation seemingly fits the notion of a ‘genomic shock’, usually denoting massive genetic and epigenetic changes that manifest immediately (or within several generations) after polyploidization (Shimizu 2022). The genomic shock includes widespread reprogramming of gene expression, activation of transposable elements (TEs), structural rearrangements, homoeologous exchange and epigenetics (Bird et al. 2018; Qiu et al. 2020), either of which has been reported in various synthetic allopolyploids. In nascent wheat synthetics prepared from *T. turgidum* and *Aegilops* sp., Ozkan et al. observed directional and reproducible sequence elimination, and an overall reduction in genome size by 4-8% occurring at the hybrid stage (Ozkan et al. 2001; Ozkan et al. 2003). Gene loss was also detected in synthetic wheat allotetraploids by Kashkush et al. (2002a), however, no sequence elimination was observed on the homoeologous group 3 by Mestiri et al. (2011) using EST markers. Dynamic changes were observed in TEs, including massive deletions and retransposition bursts (Kraitshtein et al. 2010), frequent changes in the methylation status of several TE families (Kraitshtein et al. 2010; Yaakov et al. 2010), and transcriptional activation of the Wis 2-1A retrotransposon associated with silencing or activation of adjacent genes (Kashkush et al. 2002b). A possible link between TEs and gene repression was also suggested by Li et al. (2014), who found increased density of small interfering RNAs (siRNAs) at TE-associated D-homoeologs of nascent wheat allohexaploids, which may account for their biased repression. Furthermore, Jiao et al. (2018) found differences between SSAA (*Ae. speltoides × T. urartu*) and AADD (*T. urartu × Ae. tauschii*) synthetics in their epigenetic responses (siRNA, chromatin modifications) to polyploidization.

Gene expression changes are among the most studied consequences of polyploidization, and the ‘transcriptomic shock’ has been repeatedly assessed in wheat synthetics. Prior to the availability of an annotated reference genome, changes in gene expression have been evaluated in microarray experiments, which were unable to differentiate individual homoeologous copies and could not therefore detect homoeolog expression bias. Nonetheless, the microarray studies were able to compare the total expression of a ‘gene’ (across its homoeologs and possibly also paralogs) in a synthetic to the midparent value (MPV) obtained from the parents (either by calculation or mixing of the parental RNA samples). Pumphrey et al. (2009) reported that ∼16% of wheat ‘genes’ are expressed non-additively in allohexaploid synthetics, i.e. differ significantly from the MPV. Akhunova et al. (2010) have detected an even higher proportion of non-additively expressed genes (19%) in synthetic allohexaploids; while 30.7-56.5% of ‘genes’ were found to be non-additive in the leaf and young inflorescence of allotetraploids synthesized from *T. urartu* and *Ae. longissima* (Zhang et al. 2016). Similarly high levels of non-additive expression were observed in transcriptomes of SSAA and AADD synthetics (35% and 20%, respectively) (Jiao et al. 2018). However, other microarray-based studies concluded that the vast majority of ‘genes’ in genetically stable allohexaploid synthetics is expressed additively, with only 0.7-7% ‘genes’ differing from the MPV (Chague et al. 2010; Chelaifa et al. 2013).

Relatively few studies have taken advantage of the transcriptomic approach to examine the homoeolog expression bias in nascent wheat allohexaploids. Hao et al. (2017) reported that the D-subgenome is massively affected by the transcriptomic shock, with down-regulation being the dominant change (32.6% of D-homoeologs downregulated), while changes on A and B are infrequent. Hao et al. (2017) have also established that the transcriptomic shock is not caused by the whole-genome duplication *per se*, but rather occurs at the allotriploid stage after the interploidy cross between *T. turgidum* and *Ae. tauschii*, and the transcriptional remodelling is partially reversed in the first allohexaploid generation. Ramirez-Gonzales et al. (2018) have analyzed this dataset further and found no relationship between the presence of TEs in promoter regions and altered expression patterns between homoeologs in dominant and suppressed triads.

The above overview of the published literature demonstrates little consistency of results regarding the consequences of the genomic shock in nascent wheat allopolyploids. This lack of consistency pertains to the occurrence of sequence elimination, involvement of TEs in de-regulation of nearby genes, the extent of the transcriptomic remodelling, as well as the dominant pattern of gene expression changes. Due to the conflicting results, but also a lack of comprehensively sampled data, questions persist regarding the role of the parental genotypes, maternal effects, stochasticity and heritability of the changes induced in nascent polyploids, and ultimately, about the underlying molecular mechanisms of these changes and their relevance for the plasticity of bread wheat.

Here, we present a transcriptomic analysis of several allohexaploid synthetics derived from two alternative AABB parents (Langdon, Joyau) and two alternative genotypes of *Ae. tauschii*. Our comparisons include different tissues (mature leaf and developing grain), consecutive generations, and synthetics derived from reciprocal crosses. We quantified the differential expression between the parents and the synthetics considering all genes (not limiting the analysis to triads) and TE families, analysed homoeolog expression bias and the relationship between differentially expressed genes and TEs. We identified potential sources of systematic bias arising from the difficulties of inter-ploidy comparisons involving recently-diverged species, and we describe our mitigation strategy in the Supplementary Note. Finally, we provide an updated and over-arching view on the transcriptomic shock in wheat synthetics and discuss its relevance to wheat breeding.

## Materials and Methods

### Plant material

Seeds of all genotypes used in this study were obtained from the Centre de Ressources Biologiques Céréales à paille (Small grain cereals Biological Resources Centre), INRAE, Clermont Ferrand, France. These included the *T. aestivum* L. cultivar ‘Recital’ (AABBDD) and the genotypes used for the production of the nascent polyploids - tetraploid *T. turgidum* subsp. *durum* cultivars ‘Langdon’ and ‘Joyau’ (AABB) and two diploid genotypes of *Aegilops tauschii* ‘109’ (subsp. *strangulata*) and ‘87’ (subsp. *tauschii*). All plants were vernalised for eight weeks at 4ºC (8h day / 16h night illumination regime). Four different crosses were prepared: ♀Langdon × ♂*Ae*.*tauschii*109; ♀*Ae*.*tauschii*109 × ♂Langdon; ♀Joyau × ♂*Ae*.*tauschii*109; ♂Joyau × ♀*Ae*.*tauschii*87. I.e., for the cross direction analogical to bread wheat (AABB as the maternal parent), we produced synthetics originating from the same AABB parent but different DD parents; and synthetics originating from different AABB parents but the same DD parent. Additionally, for the Langdon × *Ae*.*tauschii*109 combination, we prepared synthetics from reciprocal crosses, and we sampled two different generations. The production of synthetics involved manual castration of the female parents, rescue of the hybrid embryo followed by *in vitro* regeneration, and induction of chromosome doubling by colchicine treatment - all according to standard protocols used at our research unit.

The seeds harvested after the application of colchicine represent the first polyploid generation (labelled C1). Plants developed from those seeds were considered to be the same generation (C1), while seeds of those plants were regarded as C2. I. e., when leaf tissue and developing grain from the same plant are analysed, they correspond to C1 and C2 generations, respectively. In this sense, we sampled vigorous leafs at the anthesis/grain milk stage of the C1 and C3 generation, and developing grain at the 250 degree-days (250DD) post-anthesis, corresponding to C2 and C4 generations. All sampled plants were grown in a growth chamber, alternating 16h of light at 21ºC and 8h of darkness at 15ºC. A minimum of two biological replicates (tissues from different plants representing the same genotype and generation), sometimes accompanied with technical replicates (the same biological sample subjected to two RNA extractions) were collected. Collected samples were frozen in liquid nitrogen and stored at -80º. Detailed characterization of all samples is provided in the Supplementary Table 1.

### RNA extraction and sequencing

Multiple grains/leaves per biological sample were crushed into powder using liquid nitrogen, mortar and pestle. Up to 1g of the powder was dissolved in 4.5ml of an extraction buffer (0.1M NaCl, 10mM Tris-HCl, 1mM EDTA, 1% w/v SDS). Nucleic acids were extracted twice with 3ml phenol:chlorophorm:isoamylalcohol (25:24:1), and precipitated by adding 3M sodium acetate (1/10 of the sample volume) and two volumes of absolute ethanol. After centrifugation and resuspension of the nucleic acids in RNase-free water, the samples were purified as follows. Between 50 and 75 ug of nucleic acids were treated with 6.8 units of DNase I using manufacturer’s buffers and columns (RNase-free DNase set; Qiagen) and 1g of glycogen. Eluted RNA samples were stored at -80º and shipped on dry ice to Integragen (Evri, France), where polyA libraries were prepared with the NEBNext Ultra II mRNA-Seq protocol and sequenced on a NovaSeq sequencer (Illumina) to obtain ∼60 million PE150 pairs per sample.

### Read processing and transcript quantification

Raw RNA-seq reads produced here (62 libraries) or previously (Hao et al. 2017; Sequence Read Archive IDs SRR3474187, SRR3406932, SRR3474179, SRR3474185, SRR2474176, SRR3474199, SRR3474194, SRR3474201; henceforth referred to as ‘H2017 data’) were processed with Trimmomatic (Bolger et al. 2014), removing adapters, trimming low-quality regions and retaining only paired reads with a minimum length of 36 nucleotides each. Subsequently, two different read mapping and summarization pipelines were used for all samples. The ‘STAR-pipeline’ employed the intron-aware STAR aligner (Dobin et al. 2013) to map the trimmed reads onto a merged reference consisting of the assemblies of the *durum* cultivar Svevo v2 (Maccaferri et al. 2019) and *Ae. tauschii* subsp. *strangulata* v4.0 (Luo et al. 2017), removing reads with non-canonical intron motifs and reads with >3% mismatches against the reference. The complete ABD reference was used for all samples (including the tetraploid and diploid parents; see the Supplementary Note). Reads mapping to high-confidence genes were summarized with featureCounts (Liao et al. 2014), counting individual reads except multi-mapping reads (reads having multiple equally-good mapping choices in the reference genome) and multi-overlapping reads (reads overlapping multiple annotated features). We considered this pipeline to be optimal for the analysis of high-confidence genes, since the mapping procedure accounts for introns and the reference genome used is genetically closest to our plant material. However, this pipeline cannot evaluate the transcription of TEs, since TEs are not annotated on the Svevo assembly. This shortcoming was compensated by the second ‘bowtie2-pipeline’. Bowtie/bowtie2 (Langmead and Salzberg 2012) is often used to analyse levels of TE transcription (Navarro et al. 2019; Teissandier et al. 2019; Lerat et al. 2017; Criscione et al. 2014; Yang et al. 2019; Bendall et al. 2019; Kong et al. 2019), where intron recognition is not important and multi-mapping reads can be randomly assigned to a single location. Such random reporting of multimappers in combination with the paired-read mode gives the best quantification of TE families, according to the best practices for TE analyses from high-throughput data (Teissandier et al. 2019). While STAR (recommended by Teissandier et al. 2019) and bowtie2 have almost identical mapping percentages and true positive rates, we opted for bowtie2 which has an order of magnitude lower memory requirements - an important consideration in the case of the wheat genome. Summarization of TE-originating reads on the family level means that the pipeline is unable to quantify transcription of individual TEs, but can provide accurate overall expression of TE families. We used Bowtie2 to map the trimmed reads onto the Chinese Spring assembly v1.0 (IWGSC 2018) using the --very-sensitive-local mode and only mapping properly paired reads (--no-mixed --no-discordant). Fragments (i.e. concordant read pairs) mapping to high- and low-confidence genes (annotation v1.1) and TEs (annotation v1.0) were summarized on a meta-feature level with featureCounts, counting multi-overlapping fragments fractionally. Fractional counting was chosen to avoid inflating the library size, which could affect downstream calculations (library normalization). Prior to read counting, the TE annotation file was edited to represent each individual TE as a feature and each TE family as a meta-feature (i.e., each TE’s ID was reduced to the family level, e.g. RLC_famn30), meaning that the quantification of TE transcripts was limited to the family level. Features and parameters of both pipelines are summarized in the supplementary Table S2.

### *In silico* karyotype check and data consistency checks

Euploidy of karyotypes was determined *in silico* for each polyploid sample on the basis of median transcription over large chromosomal fragments. For this purpose, raw read counts produced by the STAR-pipeline were converted to TPMs (transcript per million). Lowly expressed genes with TPM >0.01 in less than two samples were removed, analysing seed and leaf transcriptomes separately. Parental genotypes (Langdon, Joyau, *Ae. tauschii*-87, *Ae. tauschii*-109) were presumed to be euploid, and their TPMs were averaged across replicates and used as a reference. Median TPM values were calculated for windows of 200 position-ordered genes for each polyploid sample and the reference. Corresponding median values were then compared in a ratio (sample/reference), and the resulting ratios along each chromosome were plotted on graphs. Under the expectation of euploidy, the obtained ratios should be close to one. Aneuploidy or smaller scale changes (e.g., loss of larger chromosomal fragments) should be indicated by values deviating from one (∼0.5 for monosomy; ∼1.5 for trisomy). Additionally, data consistency was checked via a clustering heatmap and a principal component analysis (PCA). The PCA was computed with RPMs in edgeR. For the heatmap, a matrix of Pearson’s correlation was calculated from log-transformed RPMs (bowtie2-pipeline; increased by +0.001 to enable the transformation) for all pairwise combinations of the samples, and the data was clustered with the UPGMA algorithm.

### Differential expression analysis

Prior to the DE analysis, a correction of ‘subgenome mismatches’ (*Ae. tauschii* reads mapped to the AB chromosomes of the reference, and *T. turgidum* subsp. *durum* reads mapped to the D-chromosmes of the reference) was applied (see the Supplementary Note). Briefly, read counts of the parents were converted to RPMs and scaled by the factors 1/3 and 2/3 for *Ae. tauschii a*nd *T. turgidum* subsp. *durum*, respectively. Subgenome mismatches of one parent were then averaged across biological replicates and the averages added to the corresponding read counts of each library of the other parent.

Additionally, TE-assigned read counts were corrected in the parental samples in the Bowtie2-pipeline. Most TE-mapping reads are multimappers and in cases when a specific mapping position cannot be identified, one of the possible mapping positions is chosen randomly (which can be on any subgenome, in both parents and the synthetics), resulting in approximately equal counts of TE reads on the A, B, and D subgenomes, even for *Ae. tauschii* and *T. turgidum* subsp. *durum*. However, all TE reads in *Ae. tauschii* and *T. turgidum* subsp. *durum* originate necessarily from the D and AB subgenomes, respectively. And since the summarization of the TE reads is performed on the family level (not on the level of individual TE copies), the read counts assigned to the ‘wrong’ subgenome (the AB and D subgenomes in the case of *Ae. tauschii* and *T. turgidum* subsp. *dicoccum*, respectively) can be added to the per-family read counts of the appropriate subgenome. Accordingly, the AB-assigned TE read counts in *Ae. tauschii* were added to the D-assigned TE read counts, while 1/2 of the D-assigned TE read counts in *T. turgidum* subsp. *durum* was added to each of the A- and B-assigned TE read counts. Similarly, the TE reads mapped to the unassigned contigs of the Chinese Spring reference genome (chrUn) were added to the TE read counts of the appropriate subgenomes in both the synthetic and the parental data sets.

After the corrections, TMM normalization (Trimmed Mean of M-values; Robinson and Oshlack 2010) was performed using the calcNormFactors function in the edgeR package. Only identical genotypes were normalized together (see Supplementary Table 1). E.g., the D-subgenomes from the synthetic samples Lx109-C2, 109xL-C2 and 109xL-C4 were normalized together with the D parent *Ae. tauschii*-109; and a separate normalization was performed for the AB-subgenome of these synthetic samples together with the AB-parent Langdon. After the normalization, genes with zero read counts across all samples were removed. Identification of DEGs between the parents and the synthetics was performed in the edgR R package, using the GLM implementation (Robinson et al., 2010). The p-values of differential expression were adjusted by the Benjamini-Hochberg method, and genes with FDR < 0.01 and fold change difference > 3 were considered to be differentially expressed.

Analysis of gene ontology (GO) was performed on the Triticeae-Gene Tribe website (http://wheat.cau.edu.cn/TGT/; Chen et al. 2020). In each GO analysis, a customized gene background was used, consisting of expressed genes (non-zero total read count) in the relevant combination of samples. A Benjamini-Hochberg correction was used on the obtained p-values, and terms with FDR <0.01 were considered significantly enriched.

## Results

### Descriptive statistics of the transcriptomes

We obtained 66 - 124 million read pairs (2×150bp) per RNA-Seq library, totalling 1.37 Tb of data (Supplementary Table 3). The read mapping and summarization, together with the downstream analyses, were performed with two different pipelines in parallel. The STAR-pipeline used the Svevo v2 and *Ae. tauschii* v4.0 assemblies as the reference genome, while the Bowtie2-pipeline used the Chinese Spring v1 assembly (for more details, see Materials and Methods and Supplementary Table 2). In the Bowtie2-pipeline that allows the analysis of TE transcription (benefiting from the comprehensive TE annotation on the Chinese Spring assembly; and also from the Bowtie2 feature that assigns multi-mapping reads to a single location, allowing family-based summarization of TE reads), 96.9%, 91.3% and 80.2% of the raw reads passed the quality-trimming, were mapped on the reference and assigned to annotated features on average, respectively. Most of the RNA-seq read pairs were assigned to high-confidence genes (86.1%; mean of all libraries), followed by low-confidence genes (8.4%) and TEs (5.5%) (Fig. 1a). Differences in TE-derived read proportions between natural (Recital) and synthetic allohexaploids were mostly insignificant, except for Jx87-C2 and 109xL-C4 (grains in both cases), where the TE-derived reads were slightly higher. Interestingly, we observed a strong and significant difference between the proportion of TE-derived reads in the grain of *Ae. tauschii* (3.5% and 4.1% of the total reads in *Ae. tauschii*-109 and *Ae. tauschii*-87, respectively) and Langdon (8.1% of the total reads). In line with expectations, the proportion of TE-derived reads in the synthetics is in between these values (4.9-6%).

**Fig. 1.**
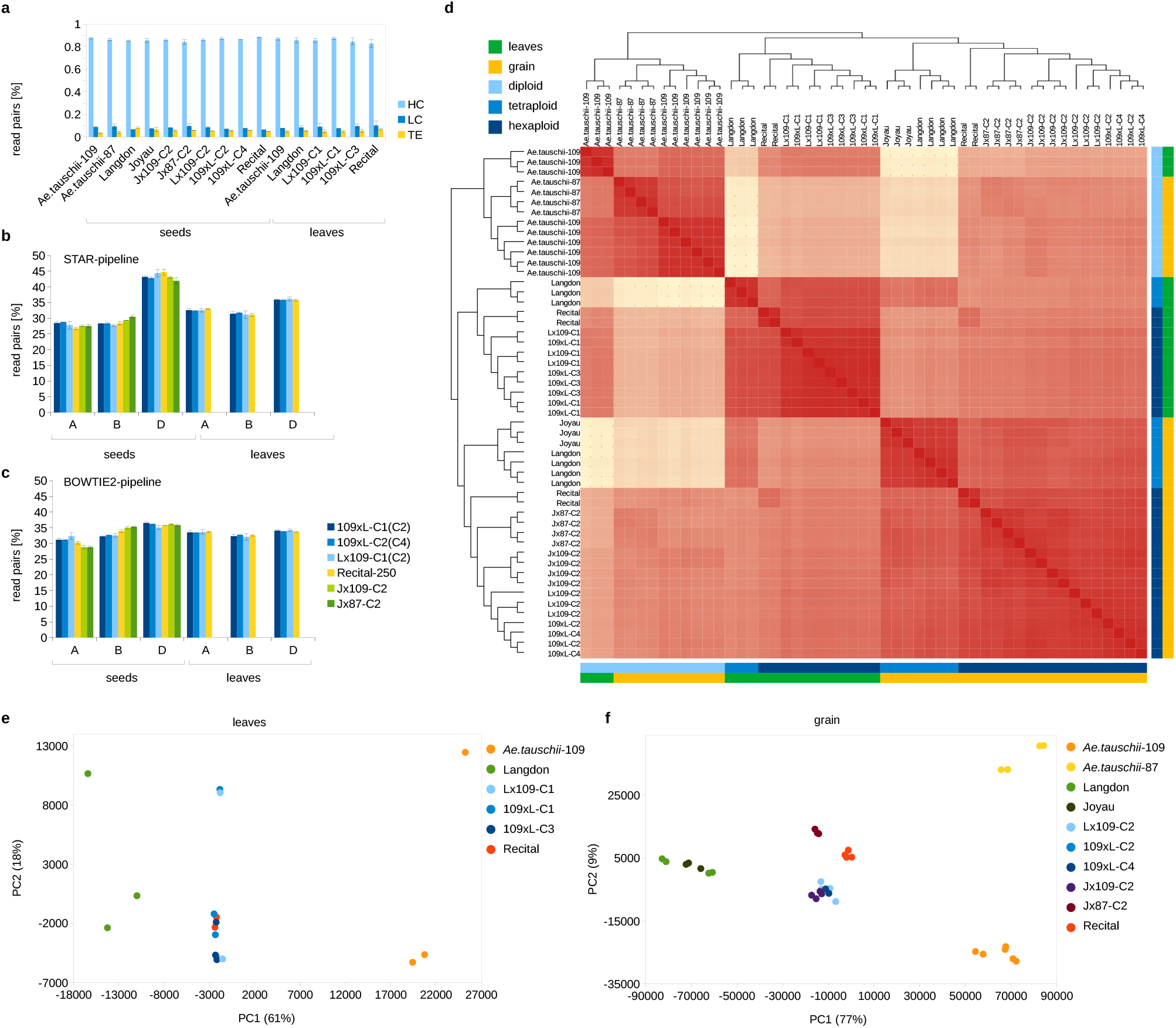
Descriptive statistics and integrity of the RNA-Seq libraries. **a**, Proportions of read pairs mapped to high-, low-confidence genes and TEs. The graph shows means of replicates, with error bars indicating standard deviation. **b**, Proportions of read pairs mapped to A, B and D subgenomes of the allohexaploid genomes. The graph shows means of replicates, with error bars indicating standard deviation for the STAR-pipeline. **c**, the same as b, but for the BOWTIE2-pipeline. **d**, A heatmap of all transcriptomes (BOWTIE2-pipeline). **e**, Top two principal components of a PCA for the leaf transcriptomes (STAR-pipeline). **f**, The same as e, but for the grain transcriptomes.

For allohexaploids, we summarized the overall contribution of the A, B and D subgenomes to the total transcription by summing up the RPMs of subgenome-assigned genes. In the STAR-pipeline, the D-subgenome contribution is apparently higher than both A and B contributions in both tissues (Fig. 1b), suggesting that D transcription dominates over the other two subgenomes. However, this could be due to systematic differences in gene annotations (e.g. different stringency for calling HC genes) in the *Ae. tauschii* v4.0 and Svevo v2 reference genomes that were used in this pipeline. In the Bowtie2-pipeline, where the Chinese Spring genome was used, the difference between D and the other two subgenomes is less obvious, though still significant in most genotypes (Fig. 1c), supporting the conclusion of a subtle D-dominance in the synthetic allohexaploids.

A heatmap of all libraries indicated good data consistency (Fig. 1d). Within the synthetic allohexaploids, reciprocal crosses and different generations originating from the same parents are sometimes intermixed, indicating high similarity of these libraries. The *in silico* karyotype check suggested that two of our transcriptomes were obtained from aneuploid samples (Supplementary Fig. S1-4). These were the leaf sample C1-ABxD-T1 monosomic for the chromosome 5B, and another leaf sample C1-DxAB-T2 monosmic for the chromosome 1B. However, the top two principal components (Fig. 1e,f) did not distinguish these monosomics, suggesting that the transcription is at least partially compensated by the present homolog. On the other hand, the PC2 of the leaf transcriptomes clearly separated four libraries that were sampled in a different year (Supplementary Table 3), indicating a technical effect that could interfere with the analysis of differential expression (DE). We therefore decided to exclude these four libraries from the DE analysis, but to keep the two monosomic samples, with a note of caution for the comparisons involving leaf transcriptomes of Lx109-C1 and 109xL-C1.

### Differential expression analysis

Our DE analysis identified hundreds of robust DEGs (fold change >3; FDR <0.01; Supplementary Fig. 2; Supplementary Table 6) between the synthetics and their parents, presumably induced by the hybridization or whole-genome duplication process. Similar DEG numbers per synthetic genotype were observed with both the BOWTIE2 and STAR pipelines (Fig. 2), and the numbers reported below refer to the former. We observed statistically significant overlaps between the sets of DEGs detected in consecutive generations (Fig. 3a-c), sets of DEGs detected in synthetics from reciprocal crosses (Fig. 3d,e), and sets of DEGs detected in different synthetic genotypes of the same generation and cross direction (Fig. 3f). Assuming these overlaps do not stem from systematic technical biases, the re-occurring DEGs testify to the non-random, and at least partially heritable and reproducible nature of post-polyploidization deregulation of gene expression. We also observed much smaller - but still significant - overlaps between the DEGs detected in the grain and leaves of the same genotype (Supplementary Fig. S5). We checked whether the low number of DEGs shared between the grain and leaves is due to large differences between the grain and leaf transcriptomes in general. This does not seem to be the case, since most genes (including the DEGs) expressed in the leaves are also expressed in the grain (Supplementary Fig. S5). Therefore, the small overlap indeed suggests that the genes de-regulated in one tissue are usually not de-regulated in another tissue, even when expressed in both.

**Fig. 2.**
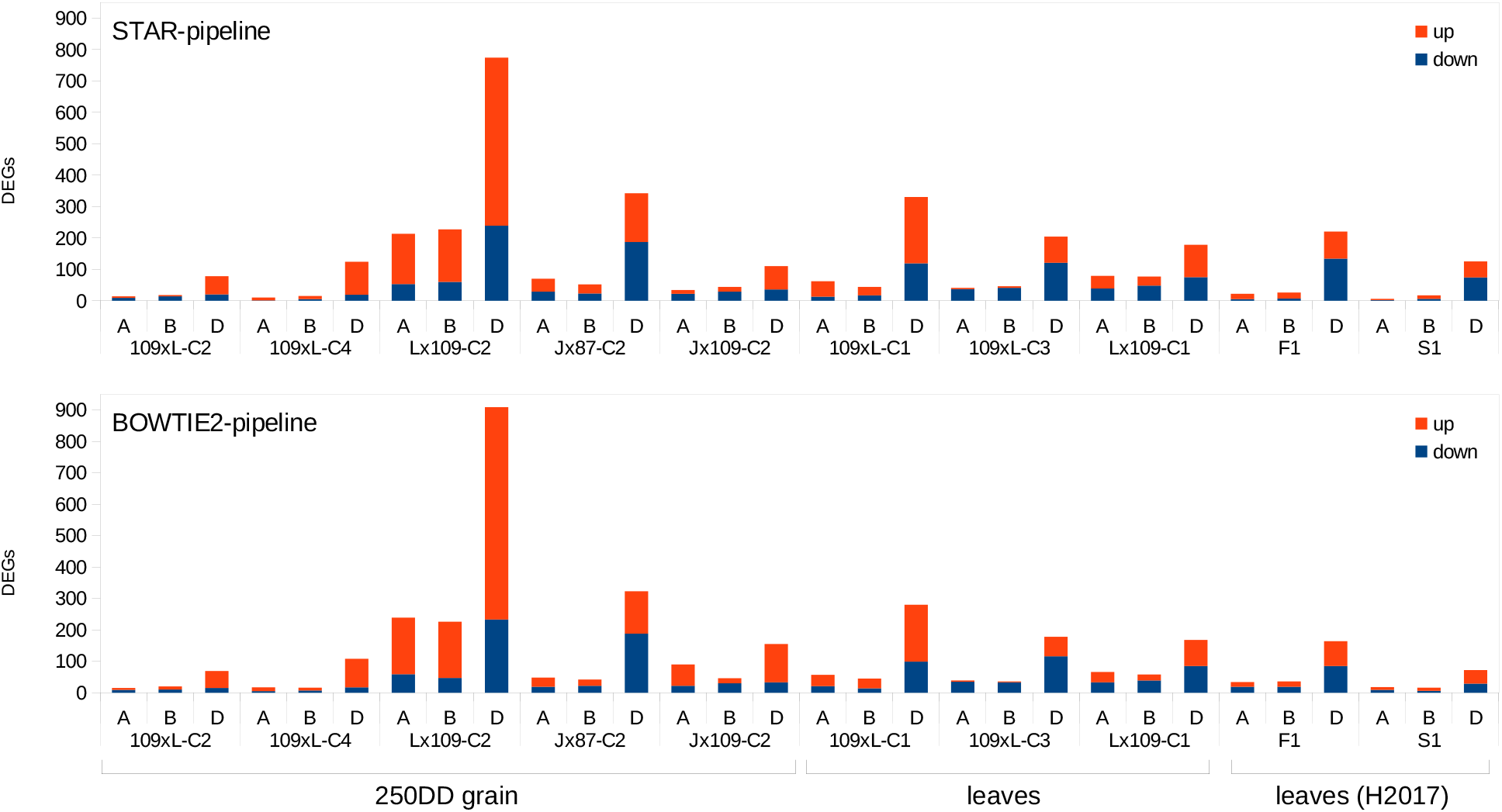
Summary of the DEGs detected by the two pipelines at the FDR <0.01 and fold change >3.

**Fig. 3.**
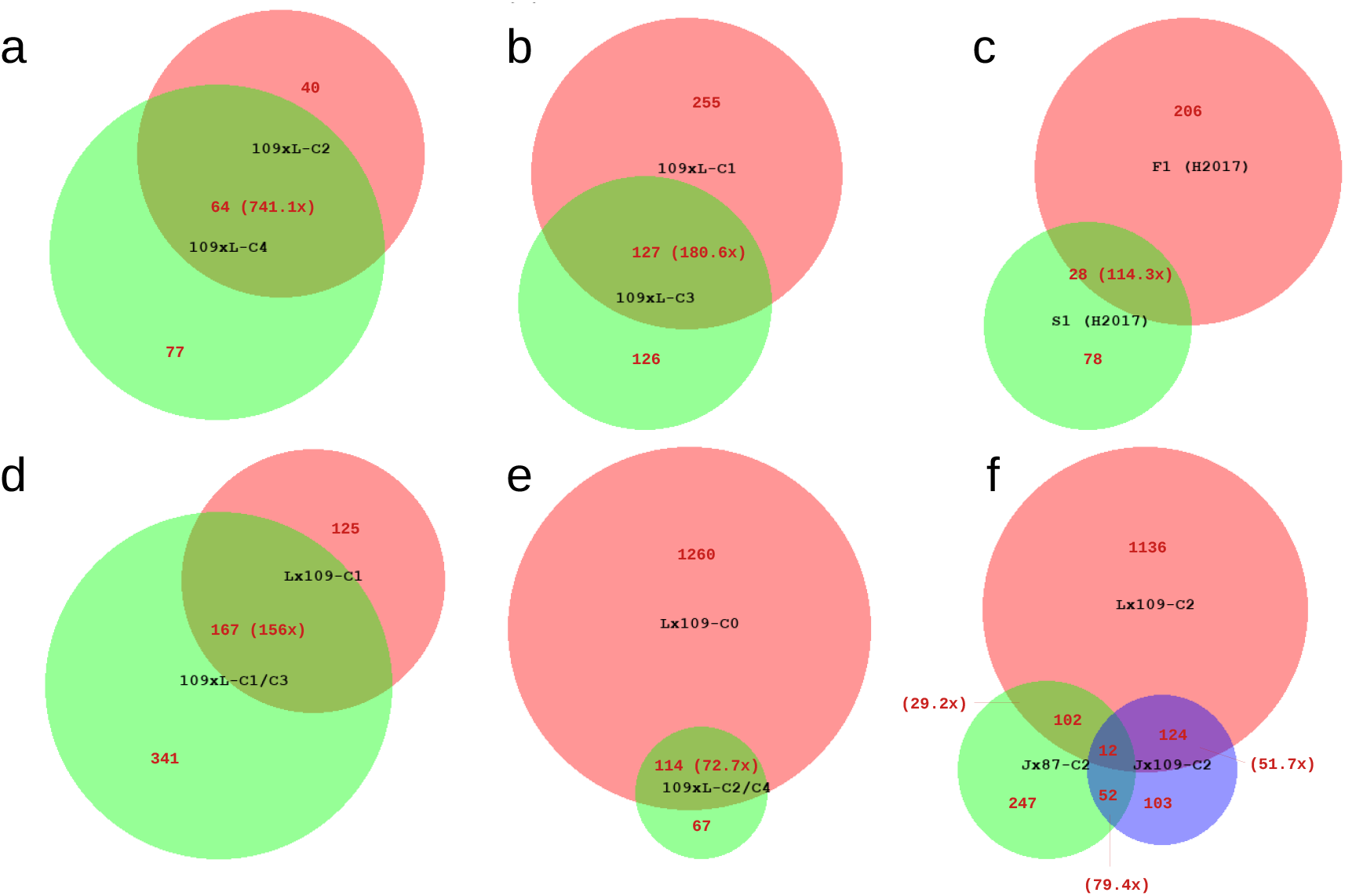
Overlaps between the sets of DEGs identified from comparisons between the parents and the synthetics. **a**, A Venn diagram showing two sets of DEGs detected in grain transcriptomes of two synthetic generations; **b**, two sets of DEGs detected in leaf transcriptomes of two synthetic generations; **c**, two sets of DEGs detected in a hybrid and a synthetic leaf transcriptome (H2017); **d**, two sets of DEGs detected in leaf transcriptomes of reciprocal synthetics; **e**, two sets of DEGs detected in grain transcriptomes of reciprocal synthetics; **f**, three sets of DEGs detected in grain transcriptomes of different synthetic genotypes. Genotypes’ ID is shown in black in the centre of the circles; DEG counts of different fractions are shown in red. The red numbers in the brackets show over-representation of shared DEGs in respect to a count expected by chance. All overlaps have high statistical significance, according to exact hypergeometric probability with normal approximation (http://www.nemates.org/MA/progs/overlap_stats.html). In these calculations, the ‘total number of genes’ was the number of genes that are expressed (non-zero read counts) in both of the corresponding DEG analyses.

Consistently across different genotypes, tissues and pipelines, higher numbers of DEGs were detected on the D subgenome. In total, we detected twice as many DEGs on the D subgenome, than we did on the A and B subgenomes together. Total DEG numbers per synthetic genotype vary considerably from 104 (109xL-C2) to 1,374 (Lx109-C2). In the grain tissue, it appears that more DEGs were yielded from crosses where durum wheat was used as the maternal parent, analogically to natural bread wheat, as opposed to crosses where the maternal parent was *Ae. tauschii* (on average, 693 and 123 DEGs, respectively). However, this was not observed in the leaves (on average, 199 and 318 DEGs, respectively). Neither was consistent the dominant direction of gene expression change. While in some synthetic genotypes and tissues, upregulation is far more frequent than downregulation (e.g., 3.05x more upregulated genes in the grain of Lx109-C2), this pattern is reversed in a different genotype (1.24-fold more downregulation in the grain of J-87-C2), or in a different tissue of the same genotype (1.16-fold more downregulated genes in leaves of Lx109-C1).

The sum of DEGs we identified in the H2017 data form a pattern that is consistent with the data that was collected by us. I.e., most of the DEGs are located on the D subgenome, without clear dominance of down- or up-regulation (Fig. 2), and with significant heritability across generations (Fig. 3c). Our results confirm the previous observation that transcriptomic changes occur already in the inter-specific hybrid, i.e. before the whole-genome duplication (Hao et al. 2017).

We performed GO analysis for each set of DEGs (i.e. separately for up- and down-regulated DEGs in each parent-synthetic comparison of DE). GO terms significantly enriched in the DEGs from either pipeline (STAR or Bowtie2) were combined, but only GO terms re-occurring across different synthetics are reported here (Supplementary Table 5). For genes downregulated in leaves, we detected significant enrichment of several GO terms related to photosynthesis and plastids. The GO terms photosynthesis (GO:0015979), light harvesting (GO:0009765, GO:0009768), chlorophyll binding (GO:0016168), protein-chromophore linkage (GO:0018298), response to light stimulus (GO:0009416), photosystem I (GO:0009522), photosystem II (GO:0009522), chloroplast thylakoid membrane (GO:0009535), plastid (GO:0009536), plastoglobules (GO:0010287) and thylakoid (GO:0009579) are all enriched within leaf DEGs downregulated in the synthetics of multiple genetic backgrounds. Some of these GO terms are also enriched among the DEGs downregulated in grain (GO:0009768, GO:0016168, GO:0018298, GO:0009522, GO:0009535, GO:0010287), indicating that polyploidization affects the same processes in various tissues. Nonetheless, several over-represented GO terms appear to be tissue-specific (Supplementary Table 5). Surprisingly, some GO terms are over-represented in both downregulated and upregulated DEGs of the same tissue (GO:0004857, GO:0004869, GO:0005576, GO:0043086, GO:0009753). E.g., GO:0004857 (enzyme inhibitor activity) is enriched among the genes downregulated in the grain of Lx109-C2 and Jx109-C2; but also among genes upregulated in the grain of Jx87-C2, Jx109-C2, Lx109-C2, 109xL-C2. Among the GO terms over-represented among upregulated DEGs are some related to stress responses. E.g., GO:0009415 (response to water), GO:0031640 (killing of cells of other organism), GO:0050832 (defense response to fungus) are terms over-represented among DEGs upregulated in the grain, while GO:0009753 (response to jasmonic acid) is over-represented among DEGs upregulated in the leaf.

### Changes in TE transcription

Although we have detected several TE families among the DEGs in the BOWTIE2-pipeline, we did not report them on the Fig. 2 and neither we include them in the Venn diagrams (Fig. 3). After closer inspection, it turned out that most of the de-regulated TE signals in the analysis of differential expression are found in TE families suffering from the ‘subgenome mismatch’ problem (see Supplementary Note). This is exposed by the RPMs in the parents, where most of the reads are assigned to the wrong subgenome (e.g., RLC_famc6 reads in *Ae. tauschii* were mapped overwhelmingly on the B subgenome rather than the D subgenome of the reference), leading to subgenomic counts that are incompatible across ploidy levels.

Moreover, many TE families show significant differences in their overall level of transcription between *Ae. tauschii* and durum wheat (Supplementary Table S4); e.g., RLG_famc7 transcripts are found in *Ae. tauschii* at RPM levels ∼500, but they are much more abundant in *T. t*. subsp. *durum* (∼7,900 RPM). Since we are unable to specifically assign TE-derived reads to the three subgenomes, the appropriate approach is to compare overall levels of transcription between the synthetics and the midparent value (MPV).

We have therefore performed an additional analysis of differential expression specifically adapted for the TEs. Unlike in the analysis of gene trascription, here we summed up the A-, B- and D-mapped RPMs (together with RPMs mapped to unassembled contigs) for each TE family. Such sum totals of the synthetics were compared to MPVs calculated for each TE class as 1/3*RPM(*Ae. tauschii*) + 2/3*RPM(*T*.*t*. subsp. *durum*), using all combination of parental replicates. Subsequently, the synthetic and midparent RPM datasets were analysed with a GLM in edgeR, using the >3 fold change and <0.01 FDR thresholds.

According to our results, the transcription level of most TE families in the synthetics is statistically similar to the MPVs (Supplementary Fig. S6). Nonetheless, several TE families appear significantly upregulated in some synthetic samples. In particular, RLC_famc4 and RLX_famc22 are upregulated in both Lx109-C2 and Jx109-C2 (grain); RLC_famc24 is upregulated in Jx87-C2 (grain); DTC_famc15 is upregulated in Lx109-C2 (grain); and DTC_famc41 is upregulated in both 109xL-C1 and Lx109-C3 (leaves). These TE families are not among those with significant parental differences (Supplementary Table S4), suggesting that a relaxation of TE silencing is indeed a likely explanation of the upregulation signals in the synthetics.

We have checked whether some TE upregulation signals could be caused by upregulation of overlapping genes and *vice versa*. We have identified only one upregulated TE family where an element overlaps with an upregulated gene on the same strand. This is a low confidence gene TraesCS5D02G548800LC (upregulated in Lx109-C2 and Jx109-C2) that is entirely located within an RLX_famc22 element. Given that the mapped reads do not follow the CDS structure of TraesCS5D02G548800LC and cover a much larger region of the TE (Fig. S7), it appears that the upregulation signal of the gene is a consequence of the upregulation of the TE family.

Additionally, we examined a possible relationship between differential expression of genes and their distance to TEs. Both up- and down-regulated genes frequently overlap with a TE, but can also be found kilobases away from the nearest upstream TE, even when strand location is considered (Supplementary Fig. S8-S15). I. e., we detected proportional amounts of significantly downregulated genes overlapping with a TE on the same and opposite strands, as well as significantly upregulated genes overlapping with a TE on the same and opposite strands. No particular TE superfamily appears to be associated with either up- or down-regulated genes (Supplementary Fig. S8-S15). Moreover, up- and down-regulation of DEGs is independent of the orientation of the closest upstream TE in all synthetics, and the same was determined for TEs located downstream of DEGs (chi-square test; alpha 0.05). I.e., the orientation of the closest TE has no relation to whether the gene is up- or down-regulated. In some synthetics, the distance to the closest TE is significantly different between up- and down-regulated DEGs, evaluating upstream and downstream TEs separately (T-test; p<0.05 in seven out of 16 comparisons). However, the pattern is not consistent across the synthetics, with TEs closer to upregulated DEGs in five cases, and to downregulated DEGs in two cases.

## Discussion

### Extent of the transcriptomic shock and the role of systematic technical biases

A transcriptomic shock, i.e. a rapid and widespread rearrangement of gene expression, is a widely-reported consequence of allopolyploidization. Some microarray studies (Pumphrey et al. 2009; Akhunova et al. 2010; Zhang et al. 2016), together with the observation of pervasive homoeologous expression bias in bread wheat (Ramirez-Gonzales et al. 2018; Mutti et al. 2018), suggest that transcriptomic shock following wheat allohexaploidization is strong. This expectation was corroborated by a transcriptomic study by Hao et al. (2017), who found that 1%, 1,2% and 25.2% of A-, B-, and D-located homoeologs, respectively, are differentially expressed (FDR <0.05) in the first allohexaploid generation compared to the parental transcriptomes (*T. turgidum* subsp. *turgidum* and *Ae. tauschii*).

In the present study, we analysed the transcriptomes in eight parents-synthetic comparisons that include different tissues (mature leaf and developing grain), synthetics produced from different parental genotypes (durum cultivars Joyau and Langdon; two genotypes of *Ae. tauschii*), synthetics produced from reciprocal crosses, and synthetics from two consecutive generations. Our DE analysis involved several key features and corrections that are important for an unbiased comparisons of transcriptomes across different ploidy levels. These include the use of the full allohexaploid reference for the read mapping of all samples (including the diploids and thetetraploids), a correction of ‘subgenome mismatches’, and separate normalization of the AB- and D-subgenomes (see Supplementary Note). Additionally, two different read-mapping and read-summarization pipelines were used, and the results of the DE analysis were compared across the two pipelines.

With a stringent fold change threshold 3 and FDR <0.01, we observed a relatively low number of DEGs following the two pipelines. We identified 110-1,224 DEGs per synthetic (STAR-pipeline; Fig. Xa), which translates to only 0.14%-1.51% of all expressed high-confidence genes (with ≥1 read detected in the compared subset of samples). Similar numbers of DEGs were identified when low-confidence genes were also considered (104-1,374 DEGs per synthetic; BOWTIE2-pipeline; Fig. Xb). Since these results are in stark contrast with the previous studies, we have included the data produced by Hao et al. (2017) in our analyses. In our pipelines, these data yielded DEGs within the same range as our original samples (Fig. Xa,b). Even after relaxing the DE thresholds (FDR <0.05, no fold change threshold, as per Hao et al. (2017)), we detected only 203 and 261 DEGs (BOWTIE2- and STAR-pipeline, respectively) in the first allohexaploid generation, compared to 4,293 in the original study. This ∼20x difference in the number of DEGs detected in the same dataset must stem from differences in the analytical pipelines. We have identified several critical points that may lead to systematic biases (Supplementary Note) and we concluded that differences in library normalizations have probably the highest impact on these two analyses. The authors of the original study compared the normalized synthetic allohexaploid libraries to the normalized libraries of *Ae. tauschii* and *T. turgidum* (M. Hao, personal communication), without taking into account the interploidy nature of such comparisons. Since the parents and the synthetics have radically different numbers of genes, transcription from a particular gene constitutes very different fractions of the total transcription in the parent and the polyploid, making direct interploidy comparison incompatible (see Supplementary Note). In contrast, we split the transcriptomes of all allohexaploids into their AB- and D-parts, and normalized the AB-parts together with the AB parent, and the D-parts with the D parent, effectively avoiding inter-ploidy comparisons. We conclude that the number of DEGs reported by Hao et al. (2017) suffers from inappropriate data normalization, and the actual fraction of genes deregulated in nascent wheat allohexaploids is ∼1%.

Library normalization is not the only potential source of systematic biases in the assessment of the transcriptomic shock. Another problem stems from the high level of similarity between the genomes involved in the polyploidization, combined with differences between the studied genotypes and the reference genome used for the read mapping. This leads to situations when reads are assigned to the wrong subgenome (here referred to as subgenome mismatches), compromising the analysis of homoeolog expression bias and DE analysis (Kuo et al. 2020; Supplementary Note). Methods developed to explicitly evaluate competing read alignments (Akama et al. 2014; Kuo et al. 2018) were found to produce much lower error rates compared to standard approaches, when used in DE analyses across ploidy levels in wheat and *Arabidopsis* (Kuo et al. 2020). This lead to suspicions that the transcriptomic response to polyploidization has been generally overestimated (Shimizu et al. 2022). Indeed, some recent studies question the very existence of the genomic shock and conclude that the changes observed in natural *Arabidopsis* and *Brachypodium* allopolyploids result from gradual post-polyploidization evolution (Gordon et al. 2020; Burns et al. 2021). Here, we placed special attention to minimize the biases related to read mapping, subgenome mismatches and data normalization (see Supplementary Note) and found that early generations of nascent wheat allopolyploids differ from their parents in a very small fraction of transcripts. This observation questions the validity of the term genomic/transcriptomic ‘shock’, at least for the polyploid system investigated here.

### Disproportionate impact on the D subgenome and the dominant direction of expression changes

Hao et al. (2017) reported that downregulation of the D-located genes in the hybrids/synthetics is by far the most frequent change in the transcription patterns, given that 95% of the detected 3,990 D-subgenome DEGs were significantly downregulated. However, downregulation was not commonly observed as the dominant pattern of non-additive expression in microarray studies, where strong (Akhunova et al. 2010) or moderate bias towards upregulation was observed (Pumphrey et al. 2009; Zhang et al. 2016; Chague et al. 2010). Here, we report a lack of consistency in the direction of significant gene expression change across tissues, genotypes and cross directions. While upregulation is more frequent than downregulation in the developing grain of most genotypes, the pattern is not consistent in leaves (Fig. Xa,b). However, we confirm the observation of Hao et al. (2017) regarding the disproportionate impact on the D genes. In all our parent-synthetic comparisons, most of the DEGs are located on the D subgenome. The overall ratio of the D-located versus A- and B-located DEGs is 1:0.27 and 1:0.22, respectively (BOWTIE2-pipeline), or 1:0.22 and 1:0.23, respectively (STAR-pipeline). This bias towards D-subgenome deregulation is unlikely to be an analytical artefact of interploidy comparisons, since our analyses were performed separately for the D and AB gene subsets (see Materials and Methods and the Supplementary Note). The cause of this bias remains unknown, but we can speculate that it relates to the fact that during the allohexaploid synthesis, the D subgenome is added to a species that is already polyploid (*T. turgidum*), with allotetraploidization estimated to have occurred ∼0.5 million years ago (Huang et al. 2002). Expression of genes that are vulnerable to deregulation due to regulatory incompatibilities of the diverged parental genomes has already been altered and stabilised in the AABB tetraploid, while such class of genes in the added D subgenome is still susceptible and therefore impacted by the genome merger.

### Induction of the transcriptomic changes and their heritability

Through the reanalysis of the data by Hao et al. (2017), we confirm that tranascriptomic changes occur already in the hybrid (i.e. allotriploid) stage. Only ∼12% of the DEGs detected in the F1 hybrid were detected as DEGs in the first allohexaploid generation, which also had less DEGs in total (Fig. Xc). This is consistent with earlier findings (Ozkan et al. 2001; Kashkush et al. 2002) reporting that some of the changes are observed at the hybrid stage, followed by additional changes in the early allohexaploid generations. Indeed, a review of genomic and gene expression consequences of allopolyploidization across several genera concluded that the majority of changes had been triggered by hybridization rather than genome doubling (Qiu et al. 2020). However, the phenomenon of nucleolar dominance is a notable exception, at least in wheat. It has been demonstrated that all parental rDNA loci are transcribed at the hybrid stage, but one of the loci is consistently silenced in the first synthetic generation of SSAA, AADD and DDAA allotetraploids (Guo and Han 2014).

It has also been proposed that genome doubling could actually ameliorate or reverse the transcriptomic changes induced by hybridization, since it has been observed that the number of deregulated genes decreases after polyploidization (Hegarty et al. 2006; Hao et al. 2017). We have also observed a marked decrease of DEGs from the hybrid (F1) to the first synthetic stage (S1; Fig. X), supporting the ‘amelioration by genome doubling’ concept. However, it is not clear whether this is a direct consequence of the doubled genome, or a simple time-dependent adjustment of gene regulation that would have occurred anyway, had the hybrids not been sterile. From a comparison of the DEG numbers between the C2 and C4 generations of 109xL allohexaploids, it appears that the transcriptome does not continue the reversal to the original parental status, i.e. cannot be adjusted further in subsequent generations (Fig. X), although this cannot be firmly concluded due to the limited number of generations sampled here.

In summary, transcriptomic changes are triggered by the hybridization step in the polyploid synthesis, with some reversals to the parental expression levels and some additional deregulation in the first polyploid generation. A notable fraction of the transcriptomic changes is heritable across subsequent polyploid generations, both in the grain and leaves.

### The role of TEs

Regarding the TE activity in general, one must distinguish between changes in TE transcription, and transpositional bursts, since the former does not necessarily lead to the latter. In this study, we focused on the changes in TE expression (summarized on the class level), and did not attempt to detect new transpositional events. While many TE classes have statistically different transcription levels in the diploid and tetraploid parents (Supplementary Table 4), very few TE classes differ significantly from the MPV. Out of ∼500 TE classes annotated in the Chinese Spring reference, only five of them were identified as differentially expressed in one or more synthetics (Supplementary Fig. SX). Notably, all of these were upregulated, and included both LTR-retrotransposons (RLC_famc4, RLX_famc22, RLC_famc24) and DNA transposons (DTC_famc15, DTC_famc41).

While no transposition bursts have been detected in nascent wheat allopolyploids (Kashkush et al. 2002; Yaakov and Kashkush, 2010; Mestiri et al. 2010), transcriptional activation of the *Wis 2-1A* retrotransposon has been reported (Kashkush et al. 2002), perhaps related to massive alterations of DNA methylation patterns observed for several TE families (Yaakov and Kashkush 2010; Kraitshtein et al. 2020). Kashkush et al. (2002) also reported that silencing of chimeric LTR-gene transcripts was associated with higher levels of antisense transcripts originating from the LTR. It has been proposed that the transcription from the LTR can generate high levels of antisense transcripts related to a neighbouring gene in the opposite orientation, which may be followed by post-transcriptional gene silencing. On the other hand, if a gene is in the same orientation as a TE located upstream, the LTR may provide an alternative promoter, which (if activated in nascent allopolyploids) can cause upregulation of the gene. However, we found that up- and down-regulation of the DEGs detected here is statistically independent from the orientation of the closest TE upstream, i.e. we cannot support the hypothesis that TE promoters are behind the upregulation of the genes in synthetics. Similarly, up- and down-regulation of DEGs is statistically independent from the orientation of the closest TE downstream, also failing to support the hypothesis that antisense readthroughs are responsible for the silencing of nearby genes. Overall, we found that only a small fraction of TE families are upregulated in nascent wheat allohexaploids, and these upregulations are not causally related to expression changes in nearby genes.

## Supplementary Figures

**Fig. S1.**
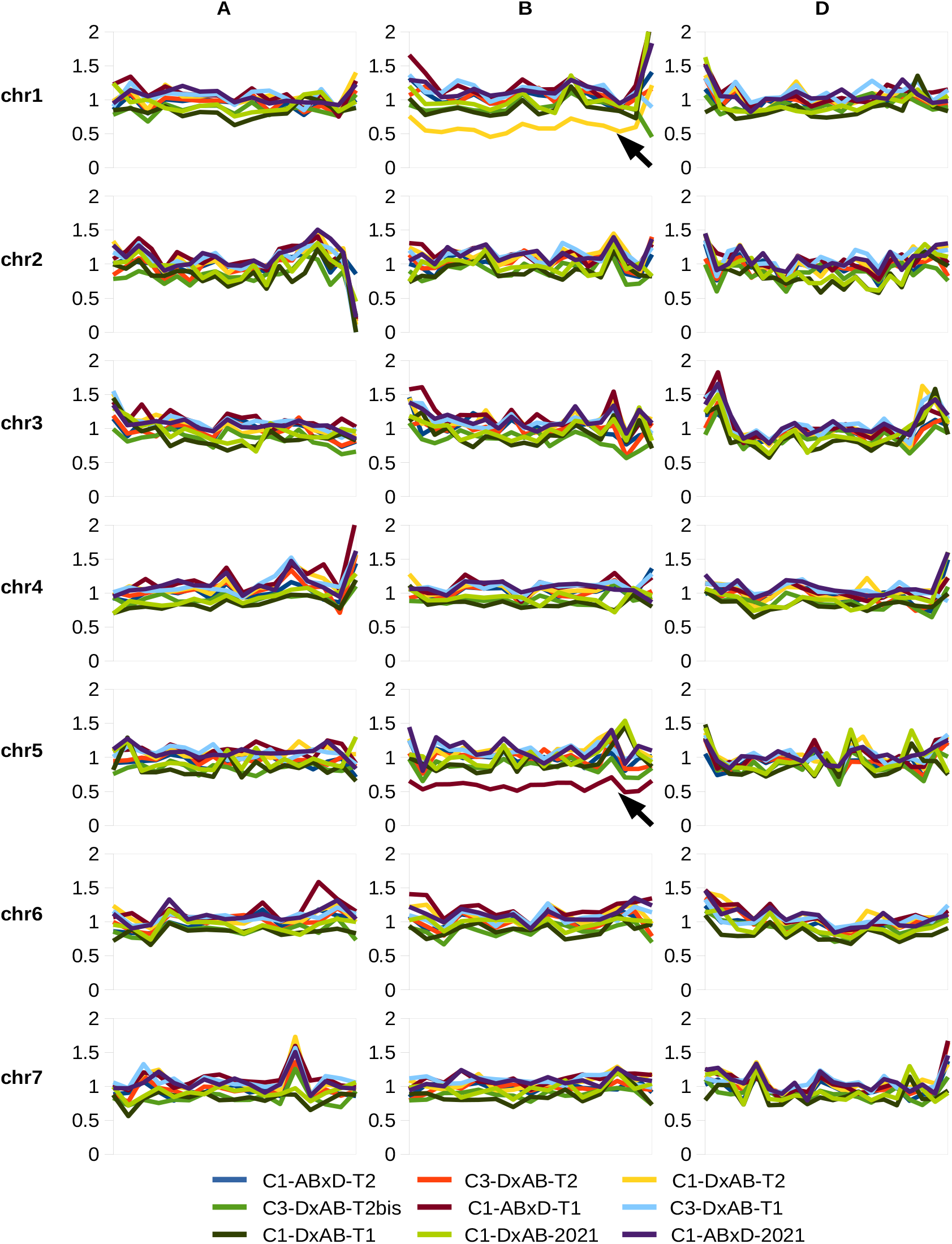
*In silico* karyotypes obtained from the leaf transcriptomes. Monosomic signals are indicated with arrows.

**Fig. S2.**
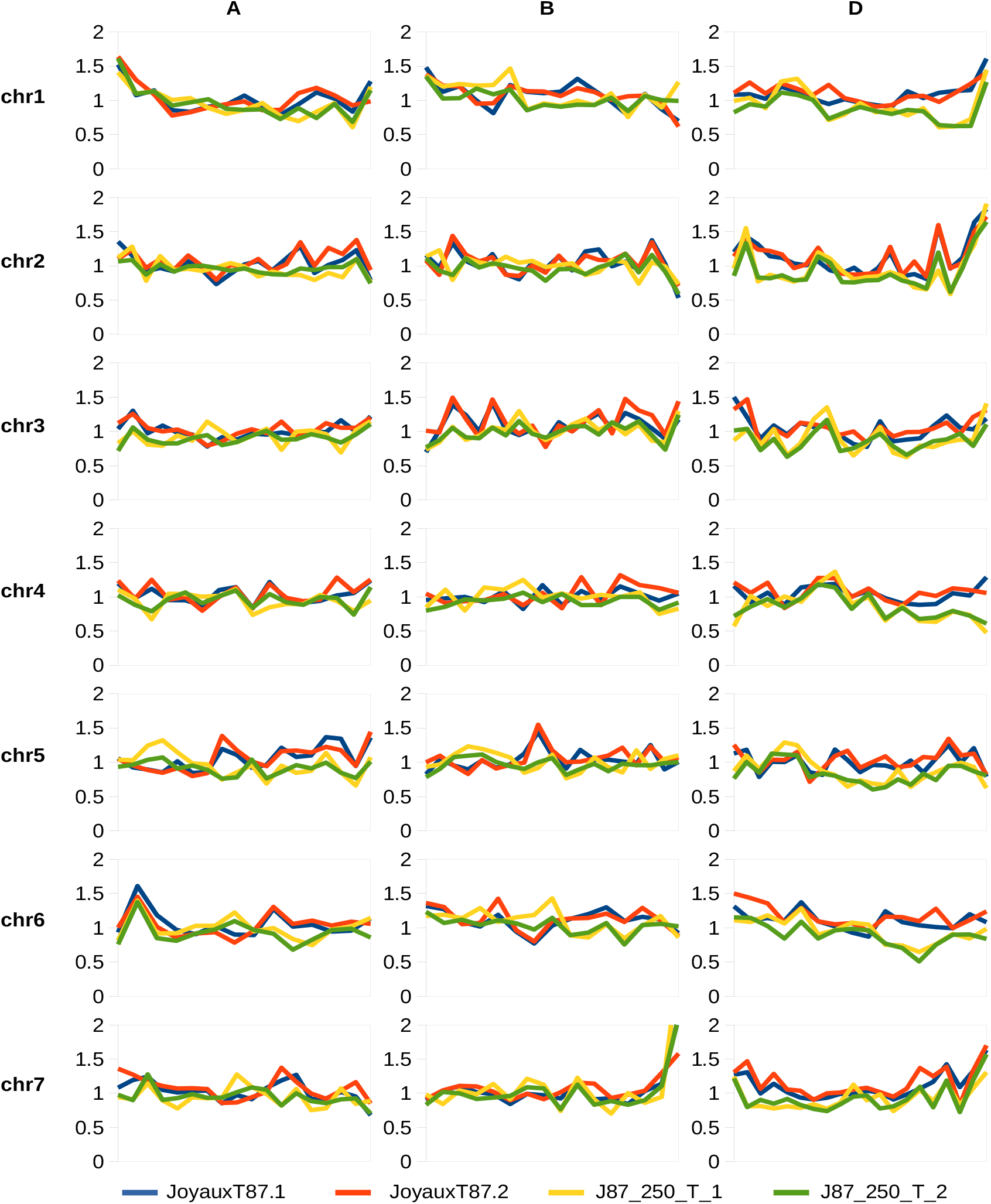
*In silico* karyotypes obtained from the grain transcriptomes of the Joyau×*Ae*.*tauschii*-87 synthetics.

**Fig. S3.**
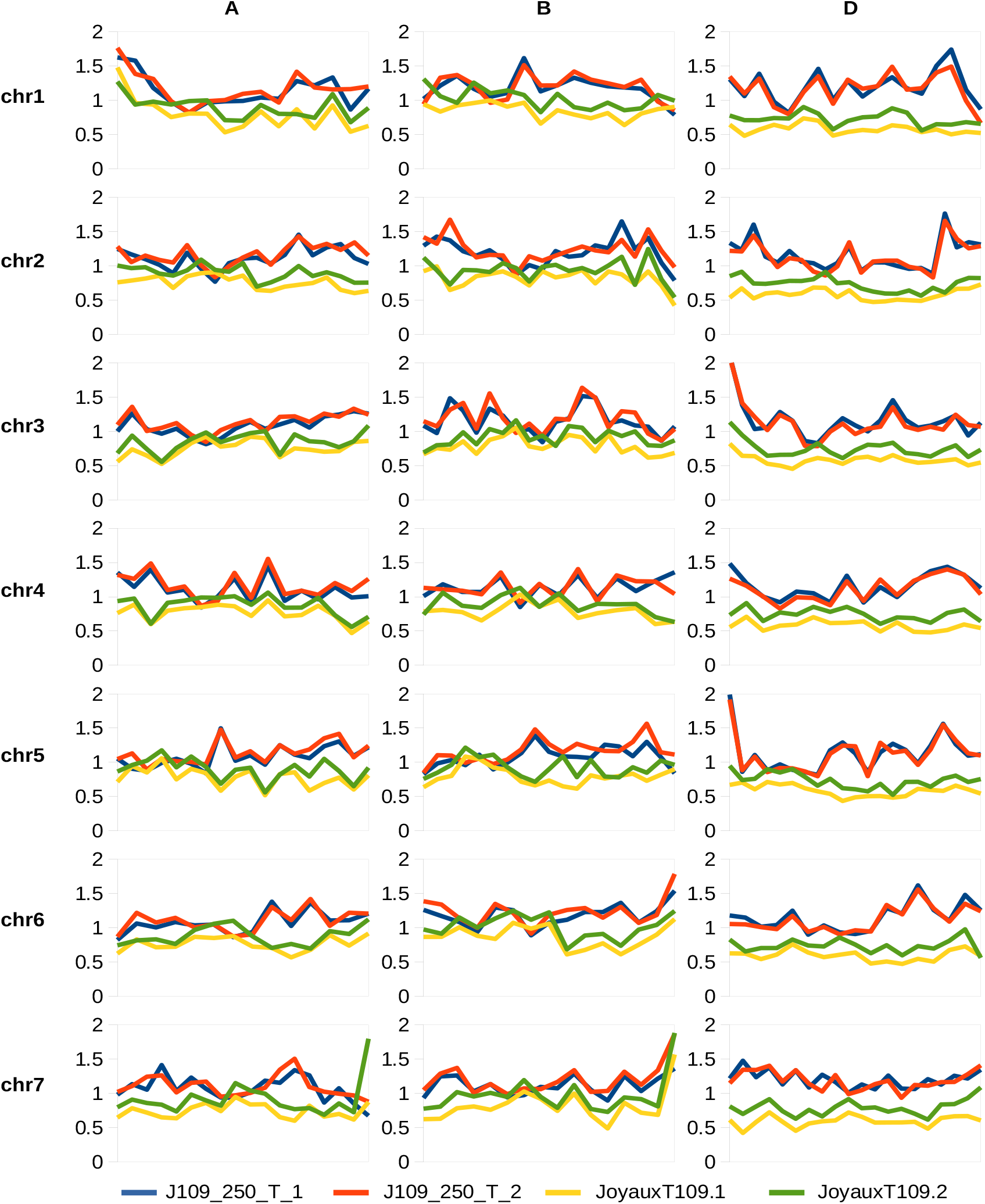
*In silico* karyotypes obtained from the grain transcriptomes of the Joyau×*Ae*.*tauschii*-87 synthetics.

**Fig. S4.**
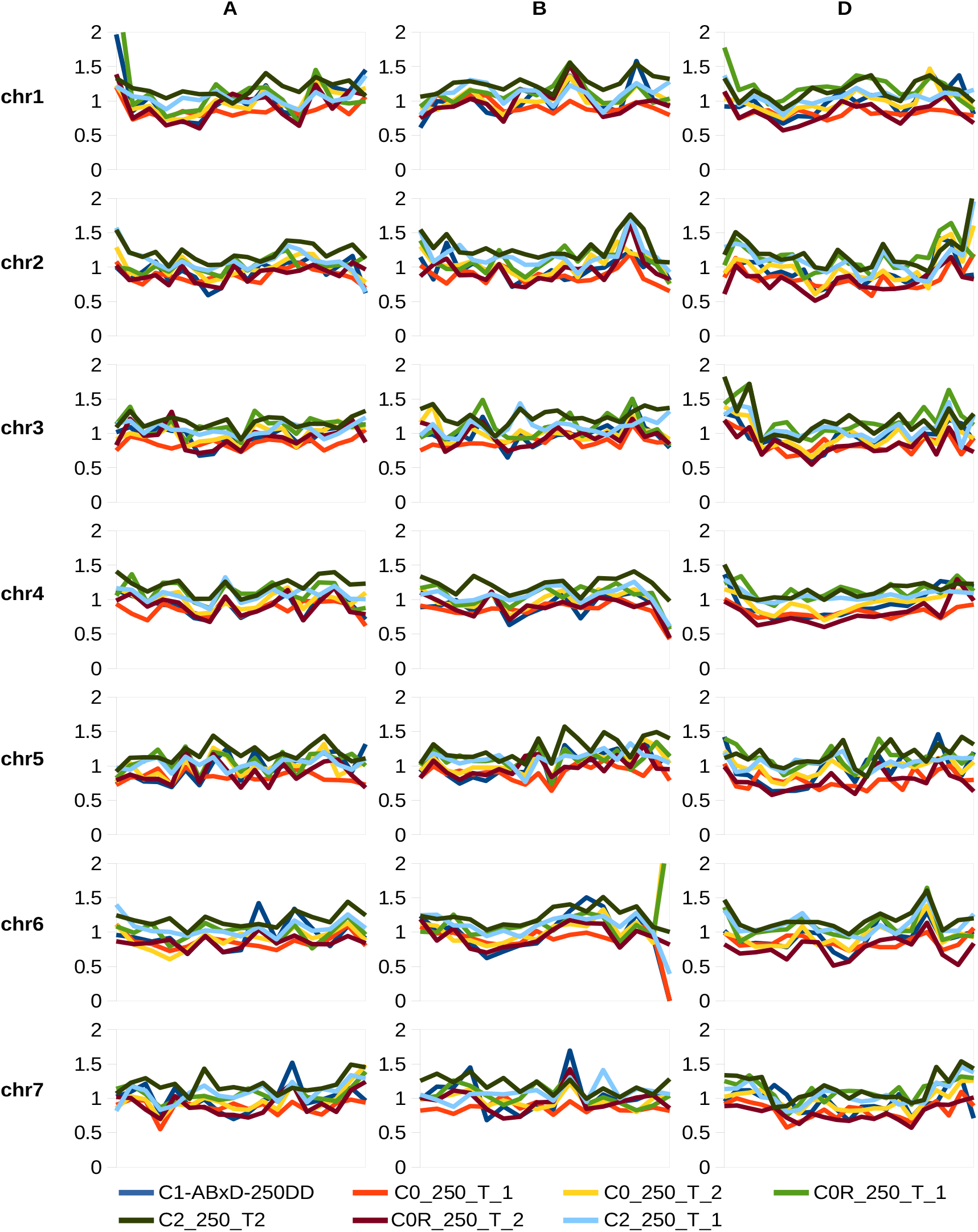
*In silico* karyotypes obtained from the grain transcriptomes of the Langdon×*Ae*.*tauschii*-109 and *Ae*.*tauschii*-109×Langdon synthetics.

**Fig. S5.**
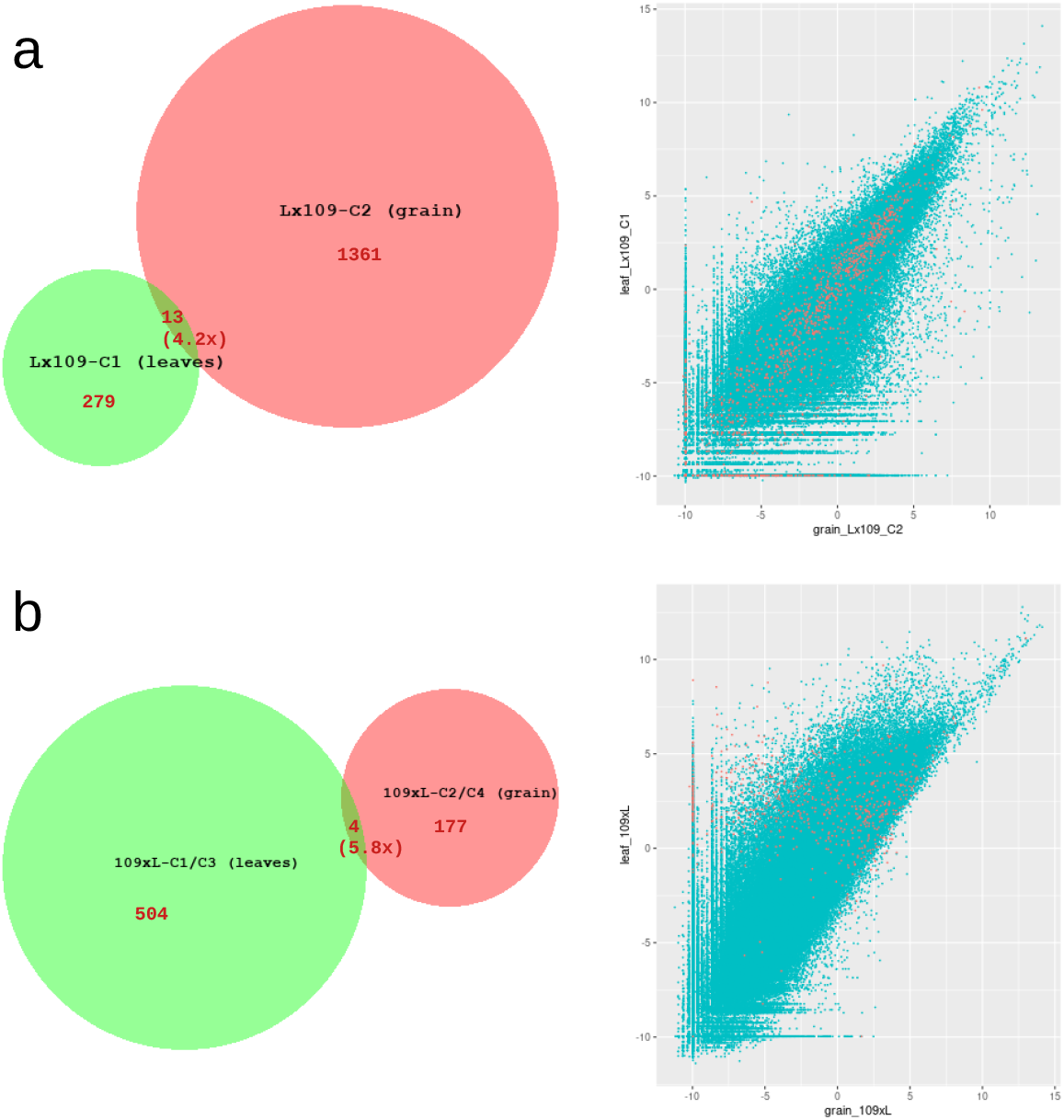
Across-tissue overlaps between the sets of DEGs identified from comparisons between the parents and the synthetics. The left panels show Venn diagrams of the DEGs (see Fig. X for more details); the right panels show a joint distribution of the expression of all genes in the grain (x-axis) and leaf (y-axis) of the given synthetic genotype. The data points are log2-transformed RPM values averaged across replicates (zeroes replaced with 0.001 before the transformation). The union of the DEGs shown on the left panels are highlighted in red on the right panels.

**Fig. S6.**
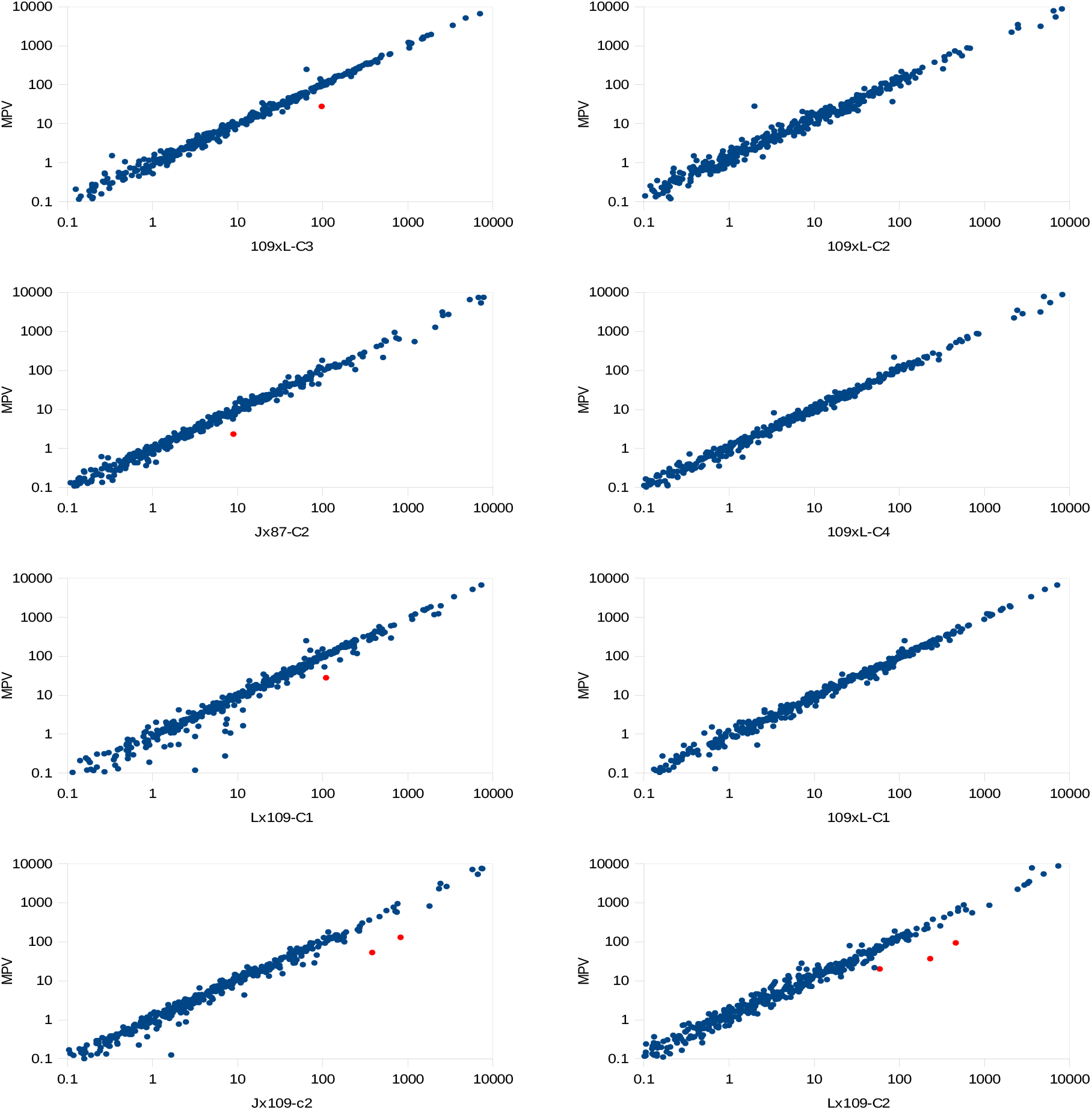
Transcription of TE classes in the synthetic wheat compared to midparent values. Transcription levels are expressed as RPMs. Each dot on the scatter plots represents a TE class; significant differences between the synthetics and MPVs are highlighted in red.

**Fig. S7.**
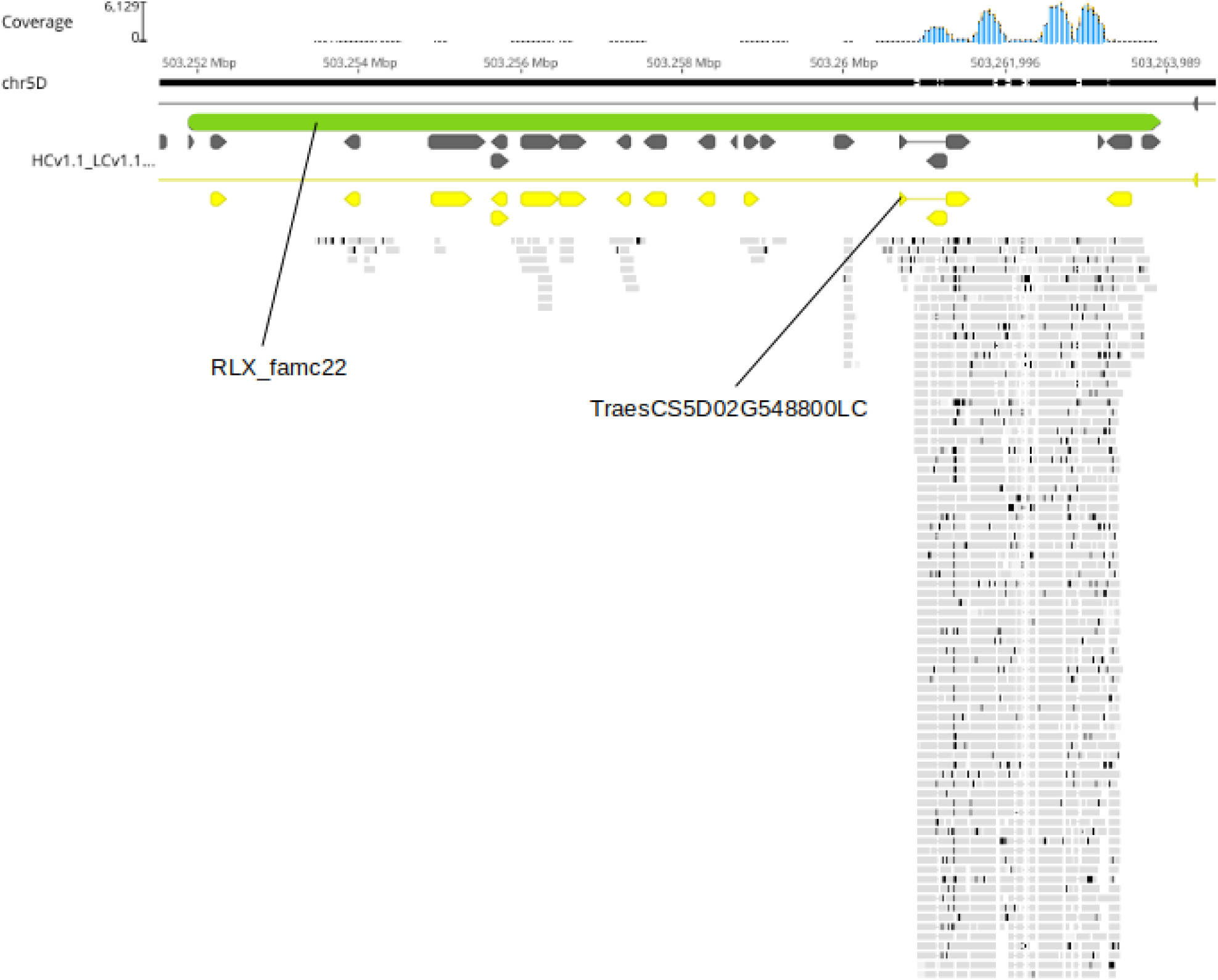
RNA-seq reads mapped to a TE element from the upregulated class RLX_famc22 overlapping with the upregulated gene TraesCS5D02G548800LC. Exons are annotated as dark-grey arrows, CDS in yellow and TEs in green; mapped RNA-seq reads are shown as light-grey bars. The figure was produced in Geneious R11 (https://www.geneious.com).

**Fig. S8.**
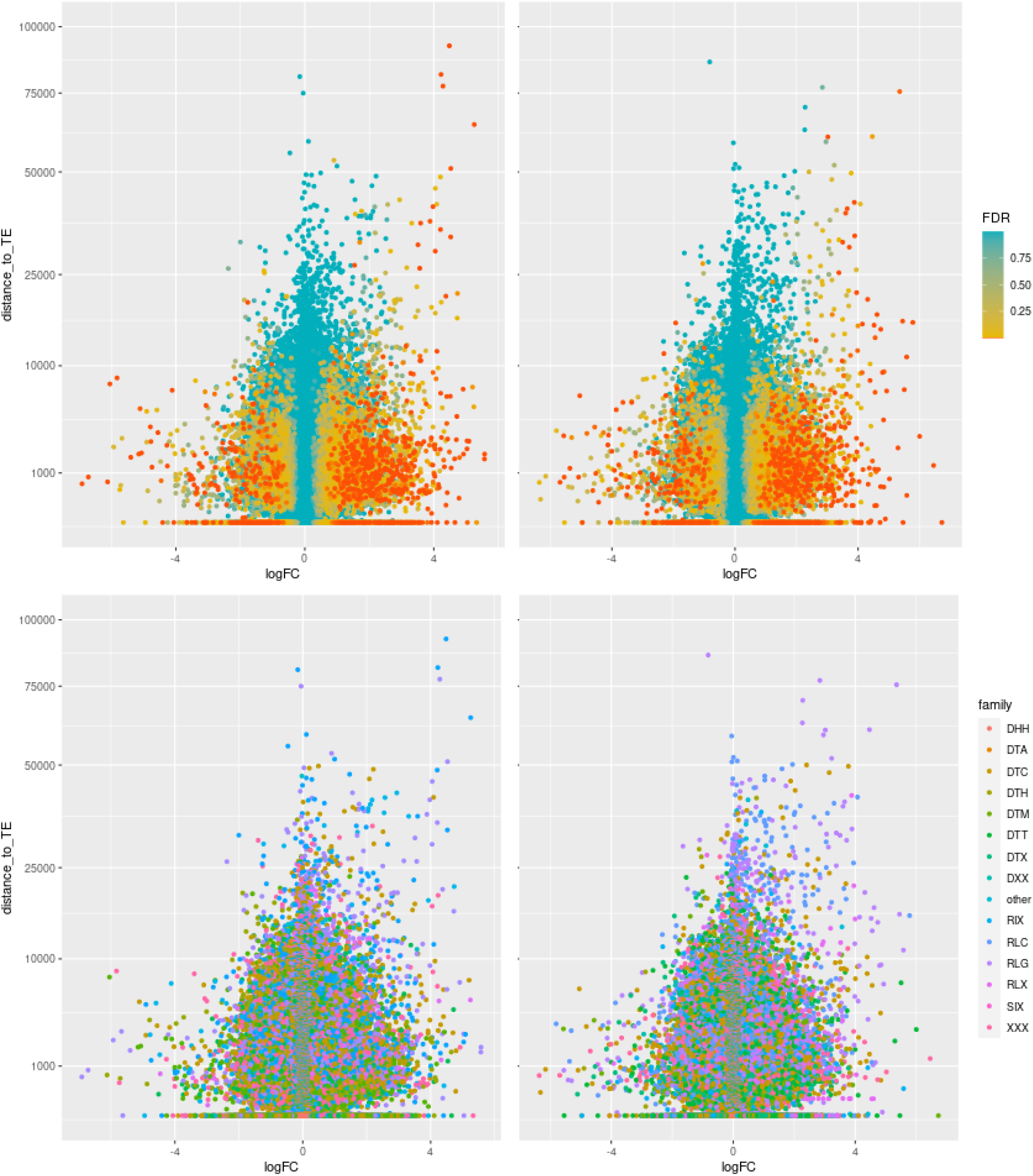
The relationship between up- and down-regulation of genes and their distance to closest TEs in Lx109-C2 (grain). The scatter plots show distance of genes to the closest upstream TE located on the same (left panels) or opposite (right panels) strands. The x-axis shows the fold change results (in respect to the parents) of the DE analysis for each gene, with the colour indicating FDR (top panels), or the superfamily of the closest upstream TE (bottom panels).

**Fig. S9.**
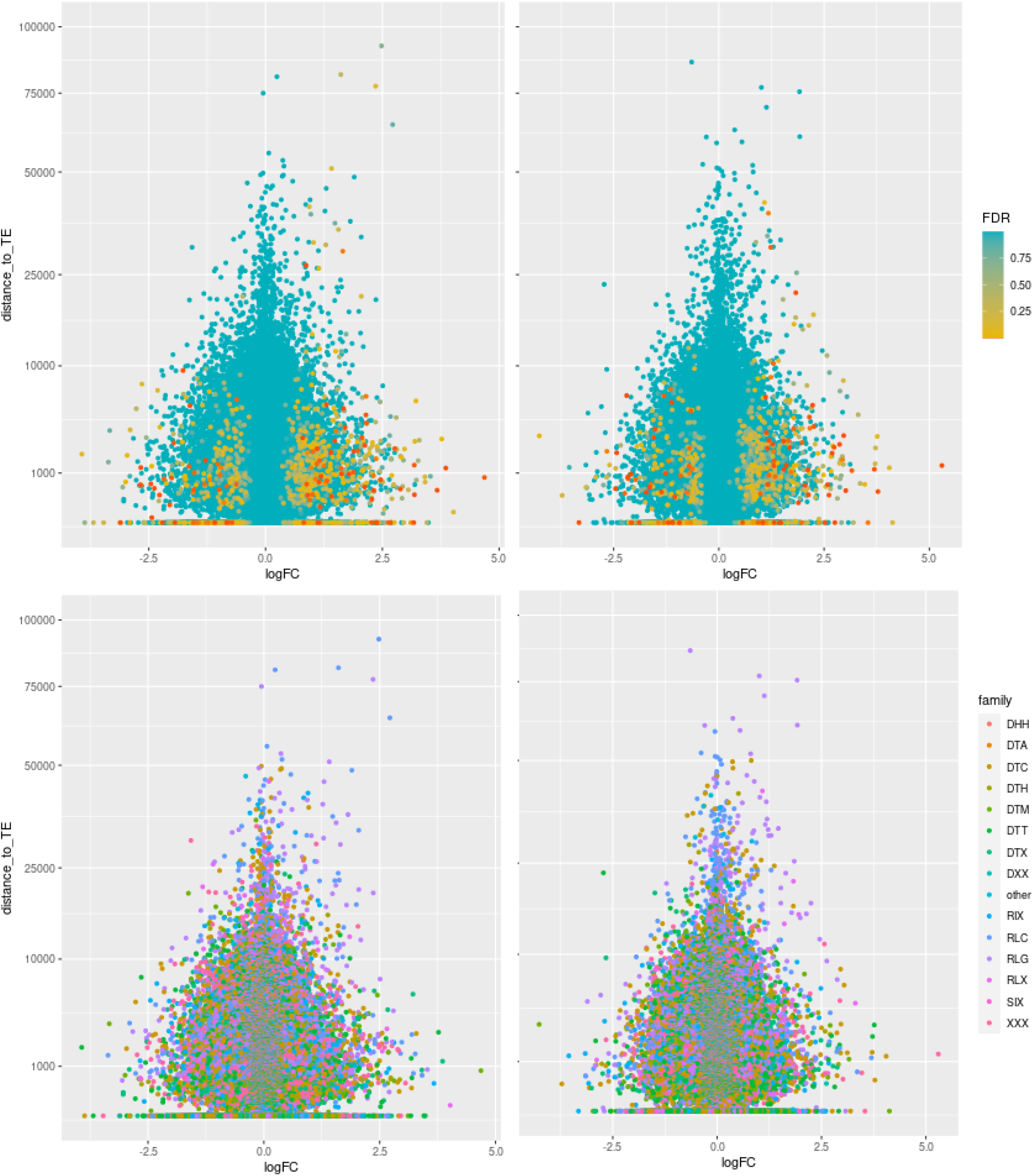
The relationship between up- and down-regulation of genes and their distance to closest TEs in 109xL-C2 (grain). See Fig. S8 for more details.

**Fig. S10.**
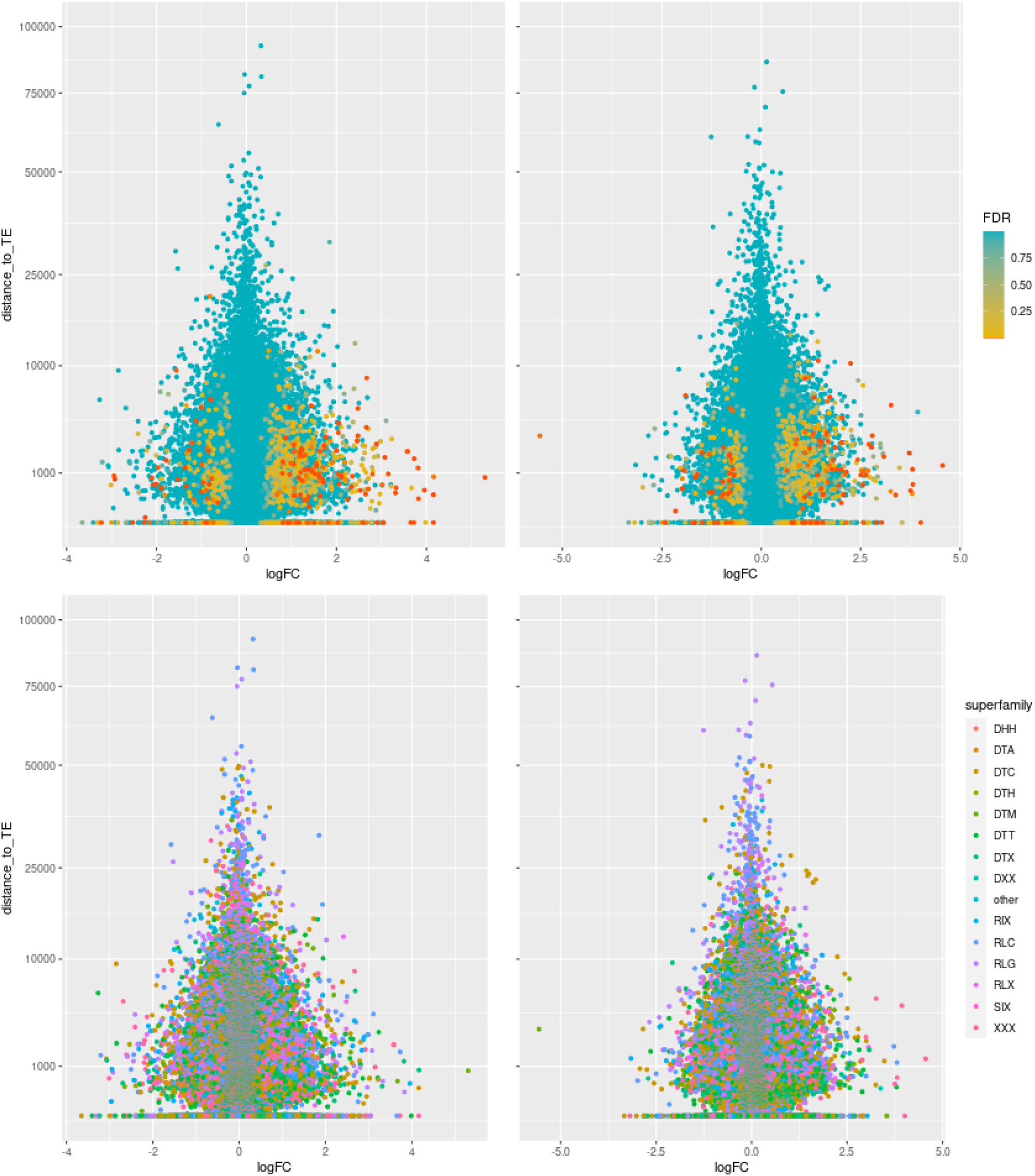
The relationship between up- and down-regulation of genes and their distance to closest TEs in 109xL-C4 (grain). See Fig. S8 for more details.

**Fig. S11.**
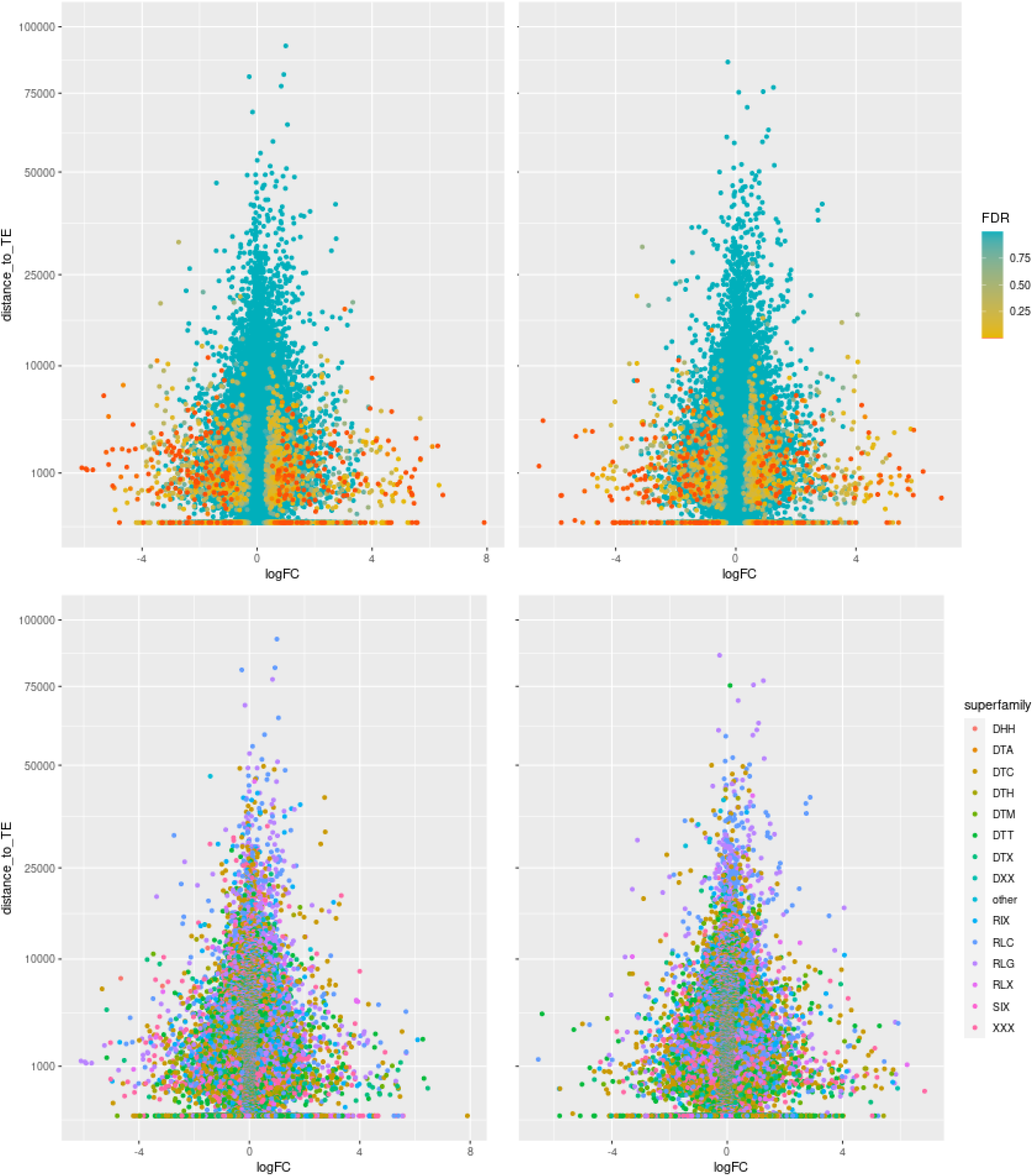
The relationship between up- and down-regulation of genes and their distance to closest TEs in Jx87-C2 (grain). See Fig. S8 for more details.

**Fig. S12.**
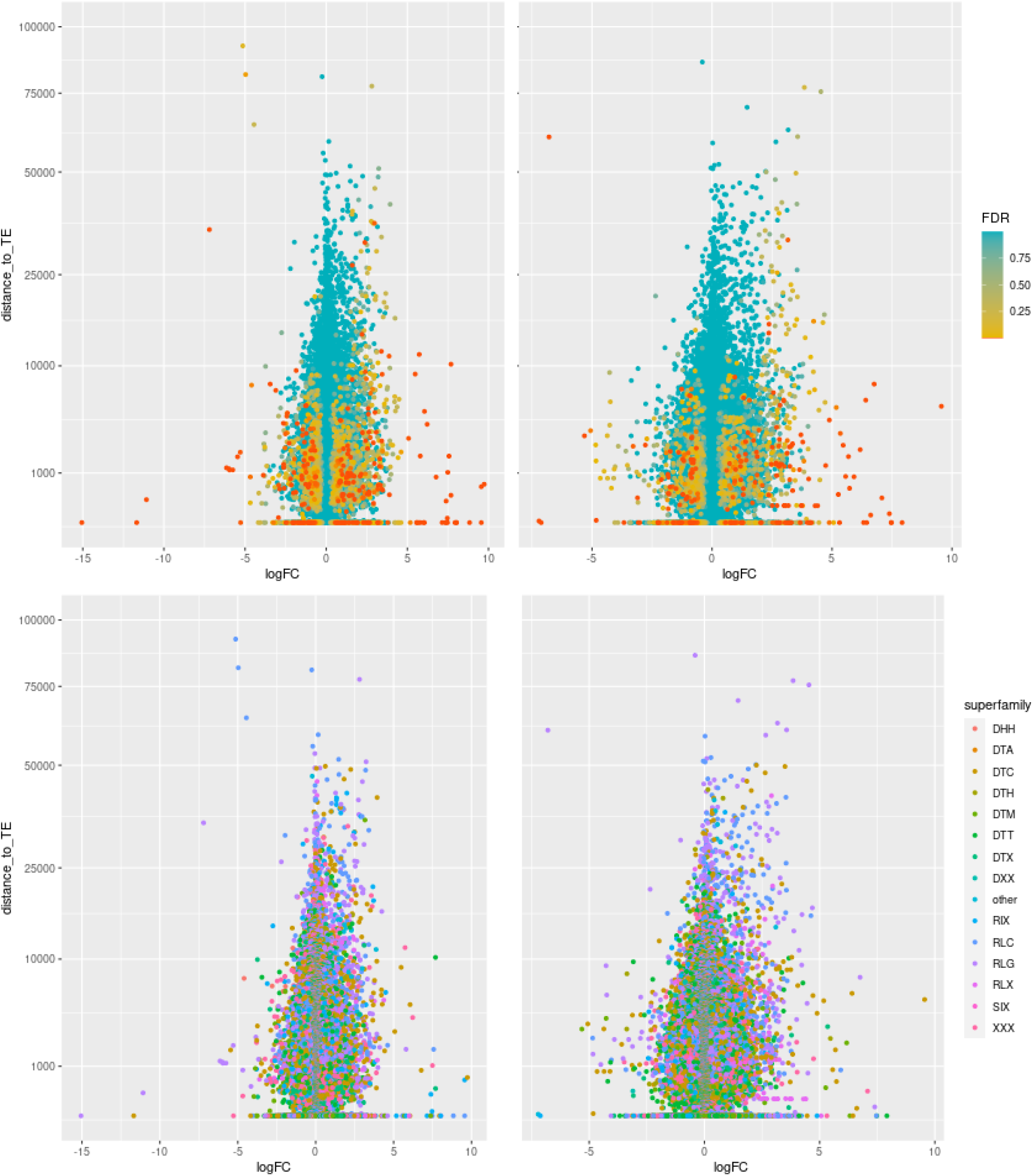
The relationship between up- and down-regulation of genes and their distance to closest TEs in Jx109-C2 (grain). See Fig. S8 for more details.

**Fig. S13.**
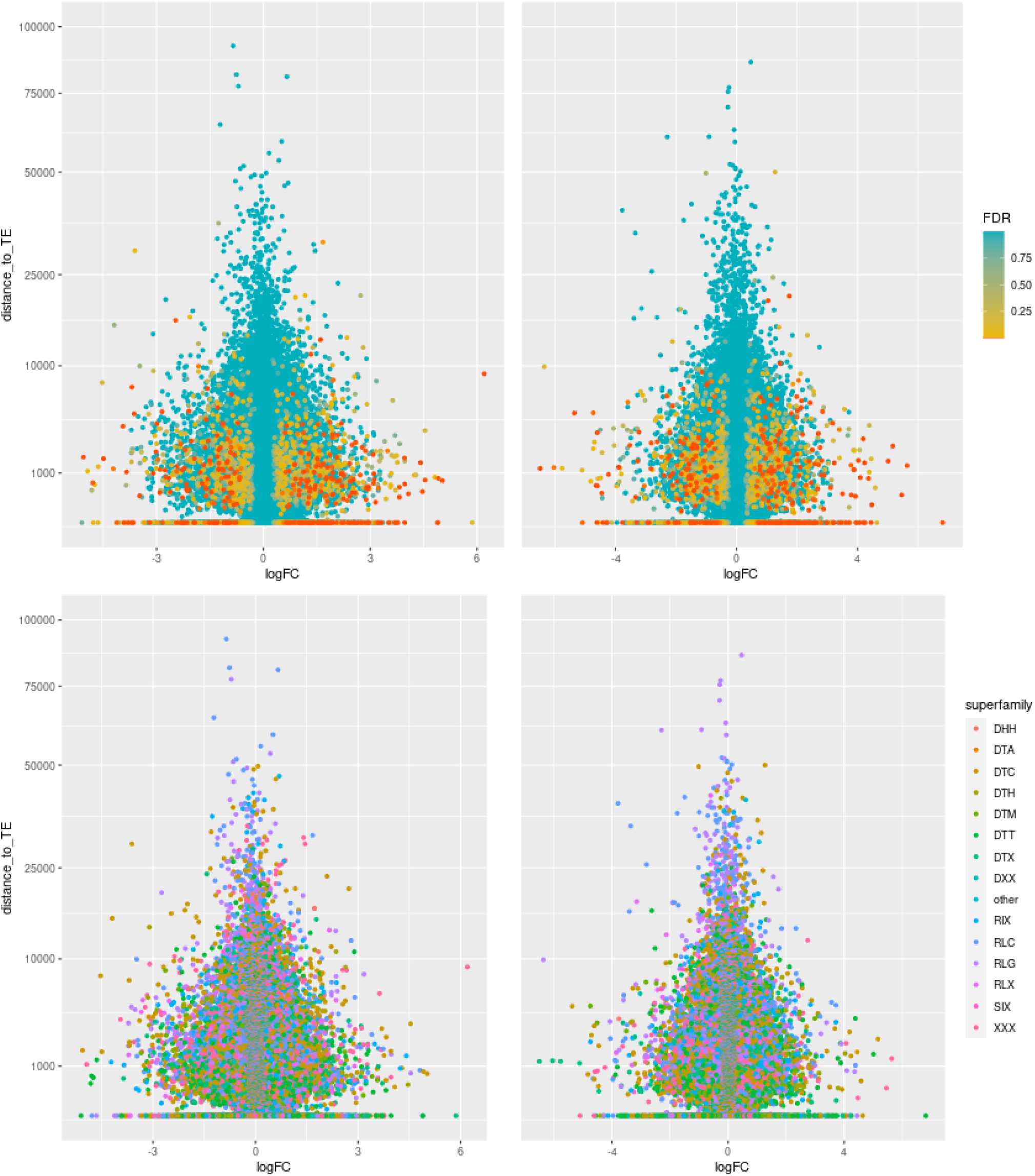
The relationship between up- and down-regulation of genes and their distance to closest TEs in 109xL-C1 (leaves). See Fig. S8 for more details.

**Fig. S14.**
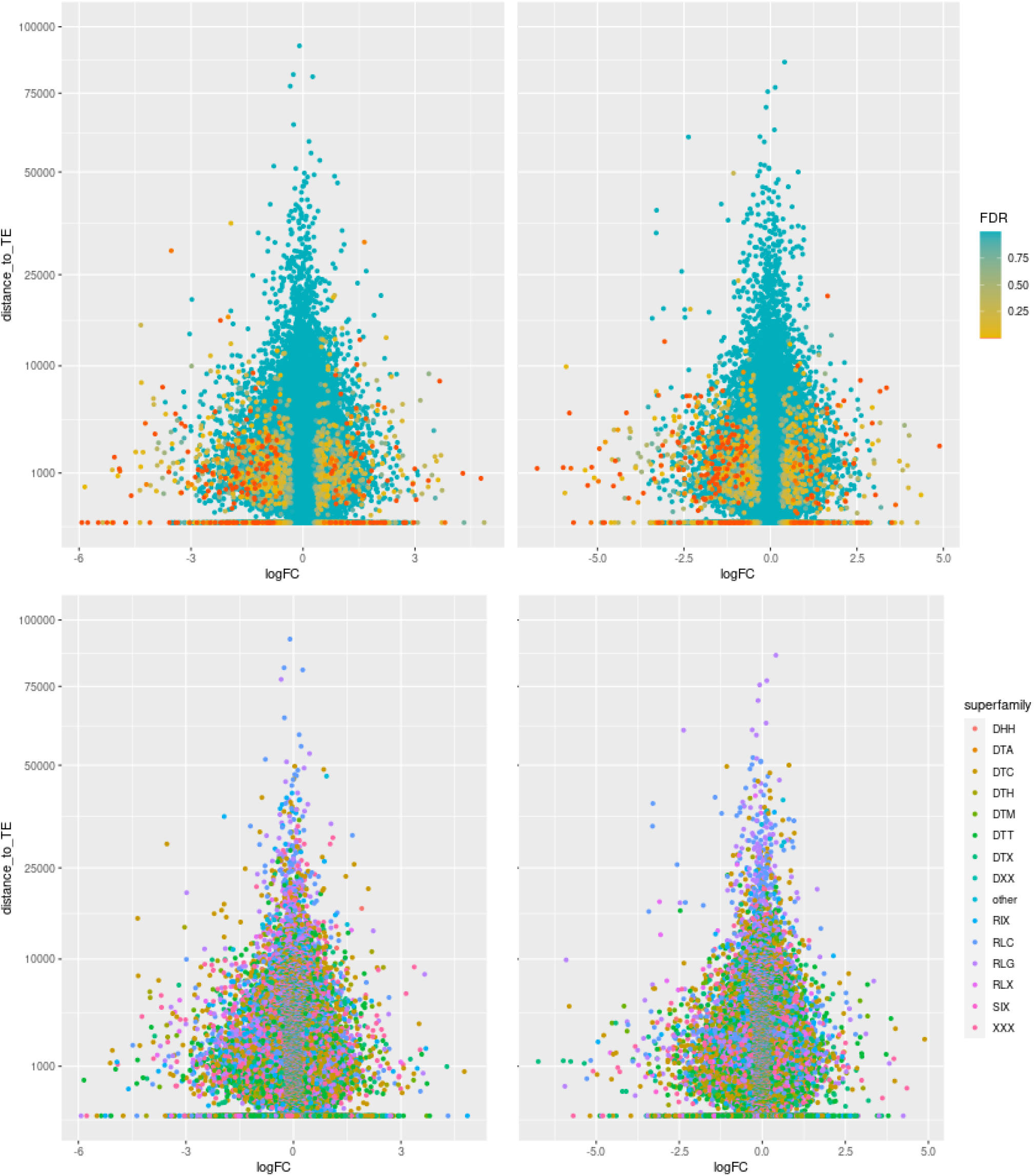
The relationship between up- and down-regulation of genes and their distance to closest TEs in 109xL-C3 (leaves). See Fig. S8 for more details.

**Fig. S15.**
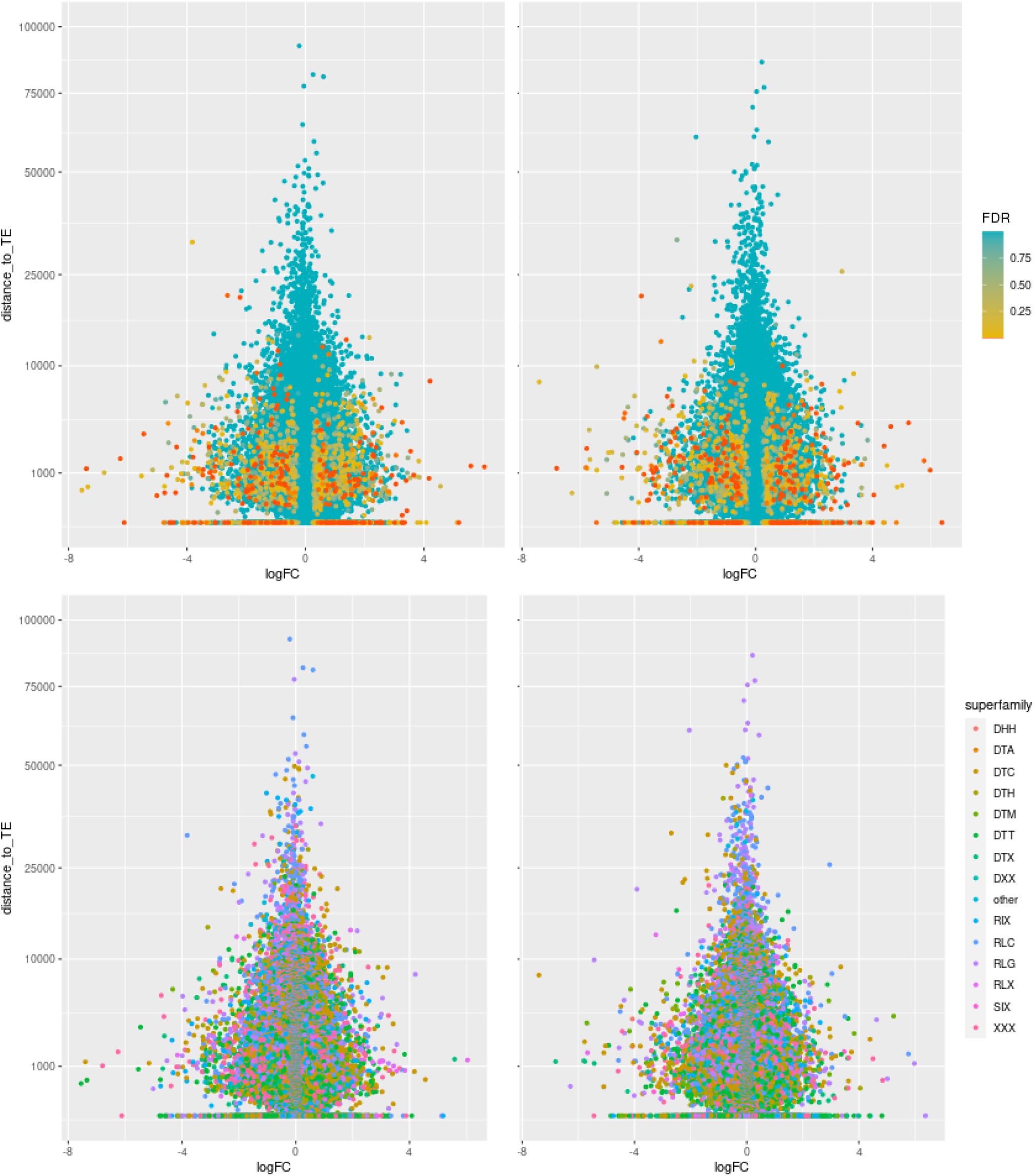
The relationship between up- and down-regulation of genes and their distance to closest TEs in Lx109-C1 (leaves). See Fig. S8 for more details.

## Notes

### Competing Interest Statement

The authors have declared no competing interest.

