## Supplementary Note for "Modest transcriptomic response to polyploidization in allohexaploid wheat synthetics"

Banouh et al.

**Supplementary Note**

**Challenges of transcriptomic analyses across different ploidy levels.**

Transcriptomic comparisons between samples of different ploidy levels are inherently difficult. While several kinds of systematic biases could conceivably lead to a large number of false positive DEGs (Differentially Expressed Genes), special care needs to be taken to prevent erroneous conclusions. Due to concerns that much of the differential expression previously reported in synthetic wheat allohexaploids could be due to the systematic biases of inter-ploidy comparisons, we took particular care to identify the possible sources of bias and minimize their impact on the results. Below, we describe several problems with demonstrable effect on the analysis of differential expression, and explain our strategy to minimize their impact.

*1. Data normalization across ploidy levels.*

The purpose of library normalization is to minimize differences between samples that arise due to systematic technical effects. E.g., when comparing two RNA-seq libraries of different sizes obtained from the same species (i.e. when different amounts of sequence data are produced), the data need to be normalized by the library size (e.g. by calculating Reads-Per-Million; RPM). However, comparing a normalized library of an ABD synthetic allohexaploid with a normalized library of its lower-ploidy parent (e.g. diploid *Ae. tauschii*) leads to a consistent signal of D-subgenome downregulation in the synthetic. This is because the D-transcription constitutes different fractions of the total transcription on the different ploidy levels (~1/3 of the allohexaploid library but 100% of the D-parent library), making the RPM counts of these two libraries incompatible. Imagine a D-located gene that generates 0.1% of all transcripts in *Ae. tauschii*, and also 0.1% of transcripts of the D subgenome of a synthetic allohexaploid (i.e. its transcription does not change after polyploidization, in respect to the total D transcription). In *Ae. tauschii*, this gene's RPM would be 1,000; however, in the synthetic (which also has the A- and B-subgenome transcripts) the RPM would be close to 333, very likely producing a significant downregulation signal. This issue cannot be resolved by more sophisticated normalization methods (e.g. TMM), since all normalization methods are designed for transcriptomes at the same ploidy levels.

One possibility for mitigating this problem is to multiply the entire parental libraries by an appropriate factor, following the logic that has been applied for interploidy comparisons of microarray data (Chagué et al. 2010; Akhunova et al. 2010). However, it is not quite clear what this factor should be (Pumphrey et al. 2009). If we imagine that all expression in an allohexaploid consists of equal parts of A-, B- and D-homoeologs, then the D-parent library should be scaled by a factor 1/3 and the AB-parent library by a factor 2/3. If we imagine that half of the expression in an allohexaploid comes from the diploid parent and the other half comes from the tetraploid parent (where, perhaps, one homoeologous copy has already been silenced), then the two parental libraries could be merged in a 1:1 ratio. Unfortunately, the real situation likely differs gene-by-gene, as observed in natural bread wheat (Ramirez-Gonzales et al. 2019), and any generalized assumption about subgenome contributions may lead to false positive signals of differential expression. Some authors therefore compare the expression of a synthetic wheat to midparent values calculated with both, 1:1 and 1:2 ratios (Akhunova et al. 2010).

We circumvent the problem of inter-ploidy normalization by restricting the comparisons on the subgenome level only. I.e., the AB parent is only normalized and compared with the AB-part of the allohexaploid libraries, and the D-parent is only normalized and compared with the D-part of the allohexaploid libraries. Consequently, this allows us to detect polyploidization-induced changes in gene expression on the subgenome level without the risk of interploidy-related biases, but this type of normalization does not allow us to evaluate contributions of individual homoeologs to the total expression of a homoeologous group. Such comparisons need to be done separately, using only the allohexaploid transcriptomes.

*2. Use of an allohexaploid genome reference for read mapping at all ploidy levels.*

It may seem natural to map the RNA-seq reads of *T. t.* subsp. *durum* onto an AB genome reference (either the Svevo reference or the AB part of the Chinese Spring reference), and the RNA-seq reads of *Ae. tauschii* to a D genome reference (either *Ae. tauschii*, or the D part of the Chinese Spring reference). However, such approach can cause problems when only uniquely mapped reads are considered in the results (Fig. 1). A read pair could have multiple equally good positions when mapping a synthetic allohexaploid on a full reference (and would be ignored during the read summarization as a multimapper), but the same read pair could have a single best position when mapping a parental genome on a partial reference (Fig. 1). Such a situation could lead to a false downregulation signal in the synthetics. To avoid this type of bias, we used the full hexaploid reference (consisting of the A, B and D subgenomes) for the read mapping of all samples, including the parental ones.

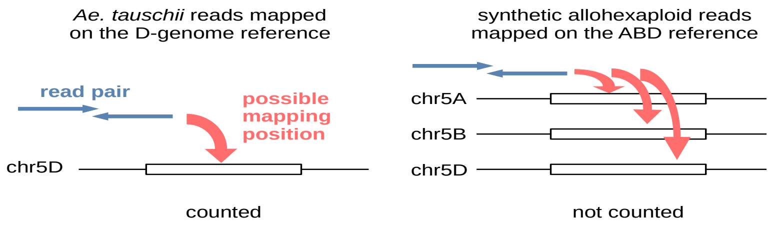


**Fig. 1.** A possible situation when a partial genome reference is used for the read mapping of the parents (left), a full hexaploid reference is used for the read mapping of the synthetics (right), and only uniquely-mapped reads are considered in the read summarization step. E.g., for a sufficiently similar homoeologous group on the chromosome 5, D-copy originating reads are not counted in the synthetic sample due to their having multiple possible mapping positions (right). However, the same reads are counted in the parent when a partial D-genome reference is used for the mapping (left), leading to a false downregulation signal in the synthetics.

*3. Correction of 'subgenome mismatches'*

When mapping lower-ploidy wheat transcriptomes on a hexaploid reference, it is commonly observed that a small fraction of uniquely mapped reads (not multimappers) are mapped to the 'wrong' subgenome. I.e., a fraction of *Ae. tauschii* reads have their best matches on the A and B chromosomes, and a fraction of *T. t.* subsp. *durum* reads have their best matches on the D chromosomes. We call such instances 'subgenome mismatches', and they likely occur due to presence-absence variation and sequence divergence between the reference and sampled genomes. In our libraries (Fig. 2), approximately 1.6% of the reads in the AB parent (Langdon, Joyau) are mapped to the D subgenome, and a similar fraction of reads in the *Ae. tauschii*-109 is mapped to either the A or the B subgenome. The fraction of subgenome mismatches is higher for *Ae. tauschii*-87 (3.1% in total), which can be explained by presumed higher divergence between this genotype and the reference sequence.

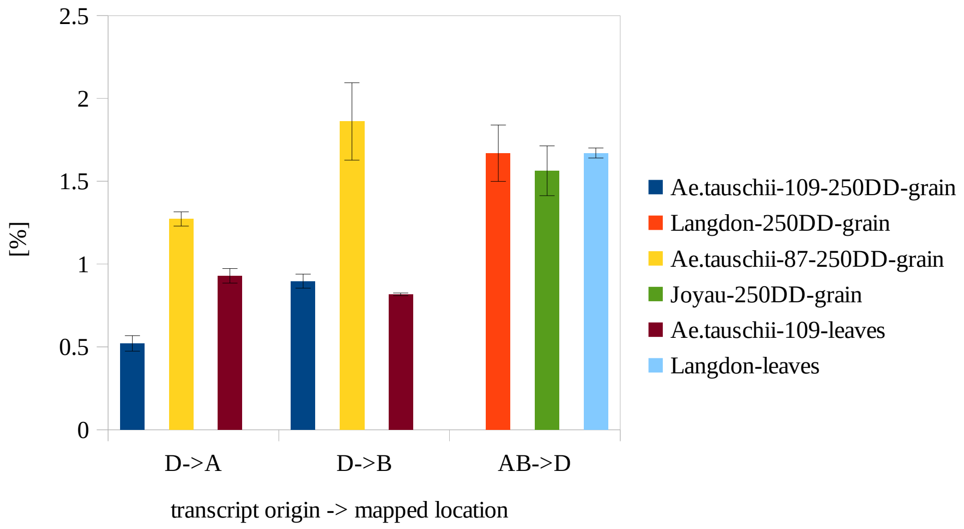


**Fig. 2.** Subgenome mismatches in the analysed libraries. Read counts were converted to RPM and the percentage of the total library mapping to the mismatched subgenome is shown, with error bars indicating standard deviation across biological replicates.

It is important to realize that subgenome mismatches are not limited to the parental genomes, where they can be easily identified. They are equally likely to occur in an allohexaploid synthesized from the same parental genomes, in which case, however, they remain invisible. Subsequently, when comparing the subgenomes of the allohexaploid with its parents (normalizing and comparing the D and AB subgenomes separately), the allohexaploid read counts will include subgenome-mismatched reads, but the read counts of the parents will not include the subgenome mismatches (Fig. 3). This could potentially lead to a false upregulation signal in the synthetics, with possibly different effects on the A, B and D subgenomes.

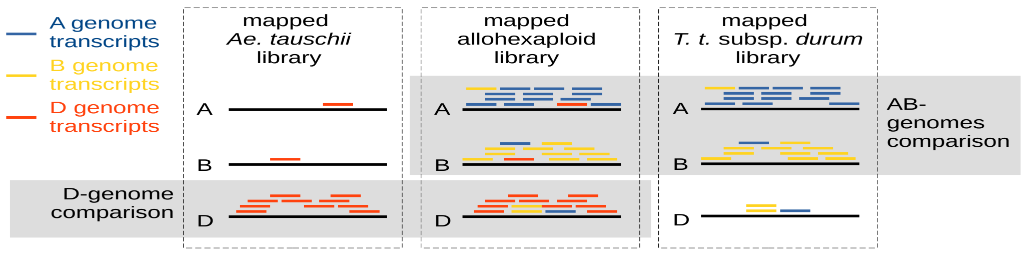


**Fig. 3.** A schematic representation of the subgenome mismatches. The black lines labelled A, B and D represent the reference genome. Note that DEG analysis is performed only between datasets within a single grey block. Because subgenome mismatches are included in the allohexaploid sample but ignored in the parental samples, false upregulation signals may occur.

Importantly, subgenome mismatches are not a consequence of using the full hexaploid reference for the mapping of parental transcriptomes, and cannot be avoided by the use of a partial reference. E.g., using the D genome as a reference for the mapping of *Ae. tauschii* could force the subgenome mismatched reads (Fig. 3; A- and B-mapped reads on the left panel) to map on the D reference (or remain unmapped), but subgenome mismatches would still occur in the allohexaploid library, potentially leading to significant differences in the parent-synthetic comparisons.

In our data, there are 1,663-2,515 genes (depending what combination of a diploid and a tetraploid genotype is used) where subgenome mismatch RPM is higher than subgenome matching RPM, i.e. genes where RPM of a D-located gene is higher in *T. t.* subsp. *durum* than it is in *Ae. tauschii*; or where RPM of an A/B-located gene is higher in *Ae. tauschii* than it is in *T. t.* subsp. *durum* (Fig. 4). If only subgenome matching counts are considered in parents, but both matching and mismatching counts are considered in the synthetics, false but statistically significant signals of upregulation can be produced for many genes. We have estimated the number of DEGs that would be detected solely due to this type of analytical artifact in our data. This was calculated under the scenario that a synthetic transcriptome equals to the one observed in the parents, but includes subgenome mismatches that are ignored in the parents. For this assessment, DEGs were identified using replicate-averaged RPMs, fold change threshold 3, and ignoring genes with RPM<1 in both the parent and the synthetic. We found 128, 114, 70 and 69 false DEGs for the Langdon×*Ae.tauschii*-87, Joyau×*Ae.tauschii*-87, Langdon×*Ae.tauschii*-109 and Joyau×*Ae.tauschii*-109 combinations, respectively.

Since these estimates of false DEGs could form a substantial fraction of the DEGs that are considered as induced by the polyploidization, it is critical to avoid this type of analytical artifact. One possibility could be to identify genes that suffer from subgenome mismatches and remove them, either before or after the DEG analysis. However, the detection of such genes is not trivial, and additionally, a gene can simultaneously be a true DEG and suffer from subgenome mismatches, potentially causing an underestimation of true DEGs. Therefore, instead of removing genes with subgenome mismatches, we attempted to correct the problem.

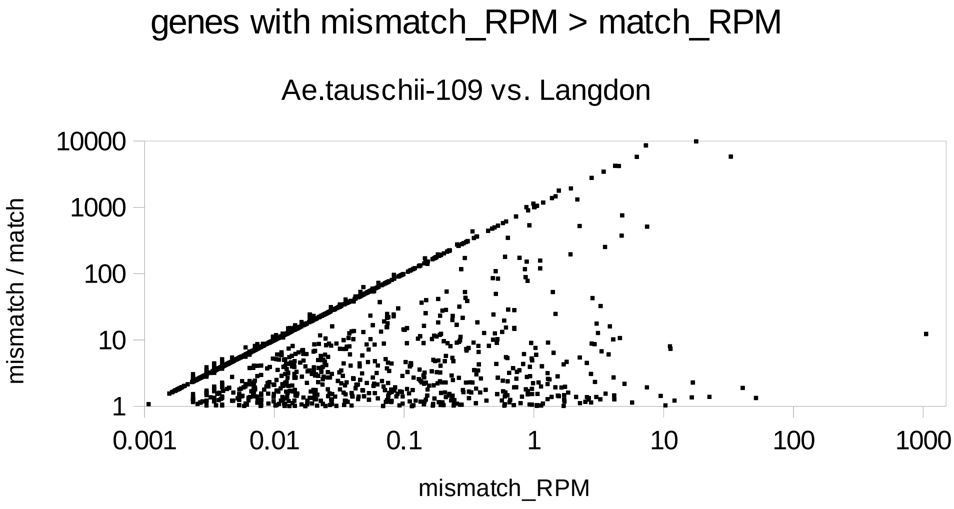


**Fig. 4.** Quantification of the subgenome mismatches. The scatter plot shows 1,663 genes where the subgenome mismatch RPM (in one parent) is higher than the subgenome matching RPM (in the other parent). This situation is observed mainly in lowly-expressed genes; however, some moderately-expressed genes are also affected.

Our correction is based on the idea that adding the subgenome mismatch read counts of one parent to the read counts of the other parent compensates the situation observed in the synthetic (see Fig. 3). Such addition needs to be performed on libraries normalized by size, and additionally, it is an inter-ploidy operation. Therefore, we first converted each parental library to RPMs, and subsequently scaled the libraries by inter-ploidy factors. I.e., we scaled the *Ae. tauschii* and *T. t.* subsp. *durum* libraries by the factors 0.3333 and 0.6667, respectively (see the part 1 of this Note). This first round of normalization allowed to perform the subgenome mismatch correction across the ploidy levels in the parents: subgenome mismatches of one parent were averaged across replicates and the averages added to the corresponding normalized counts of each replicate of the other parent. The corrected parental libraries were then used in another round of normalization, where the TMM method was applied separately for the AB- and D-subgenome, following the logic of separate normalization (see the part 1 of this Note) and allowing the DEG detection unbiased by inter-ploidy comparisons.
